# Expression of neuronal Na+ leak channel, NALCN, provides for persistent invasion of metastasizing cancer cells

**DOI:** 10.1101/2020.08.13.249169

**Authors:** Oksana Iamshanova, Dmitri Gordienko, Antoine Folcher, Alexandre Bokhobza, George Shapovalov, Dheeraj Kannancheri-Puthooru, Pascal Mariot, Laurent Allant, Emilie Desruelles, Corentin Spriet, Raquel Diez, Thibauld Oullier, Séverine Marionneau-Lambot, Lucie Brisson, Sandra Geraci, Hathaichanok Impheng, V’yacheslav Lehenkyi, Aurelien Haustrate, Adriana Mihalache, Pierre Gosset, Stéphanie Chadet, Stéphanie Retif, Maryline Laube, Julien Sobilo, Stéphanie Lerondel, Giulia Villari, Guido Serini, Alessandra Fiorio Pla, Sébastien Roger, Gaelle Fromont-Hankard, Mustafa Djamgoz, Philippe Clezardin, Arnaud Monteil, Natalia Prevarskaya

## Abstract

Cytosolic Ca2+ oscillations provide signaling input to several effector systems of the cell. These include neuronal development, migration and networking. Although similar signaling events are hijacked by highly aggressive cancer cells, the complexity of the ‘neuron-like’ remodeling in metastasis remains to be explored. Here, using a variety of in vitro and in vivo techniques we show that strongly metastatic prostate cancer cells acquire specific Na+/Ca2+ signature required for persistent invasion. We identify the ‘neuronal’ Na+ leak channel, NALCN, at the hot spots of the Ca2+ wave initiation and invadopodia formation. Mechanistically, NALCN associates functionally with plasmalemmal and mitochondrial Na+/Ca2+ exchangers, reactive oxygen species and store-operated channels to generate intracellular Ca2+ oscillations. In turn, this stimulates the activity of protooncogene Src kinase co-localized with NALCN, actin remodeling and secretion of proteolytic enzymes, thus increasing an invasive potential of the cancer cells and metastatic lesions in vivo (accessed in pre-clinical models). Overall, our findings provide new insight into the signaling pathway specific for metastatic cells where NALCN plays the role of the persistent invasion “launcher and controller”.

## Introduction

Prostate cancer (PCa) is the most common noncutaneous human malignancy and the second most lethal tumor among men, with its highest incidence in industrialized countries. Understanding the processes facilitating the progression of PCa to aggressive metastatic phenotypes and developing new therapeutic targets are necessary to improve both the survival rate and the everyday life of the patients. In most cases of PCa, metastasis but not the primary tumor *per se* is the main cause of mortality, with the bone metastases being the most likely and highly incurable complication.

To effectively escape the tumor, enter the circulation and establish secondary growth in distant organs cancer cells must develop an enhanced migration propensity. The invasion capacity gained by tumor cells is the hallmark of malignancy. To become invasive, tumor cells need to acquire traits enabling enhanced migration and proteolysis of extracellular matrix (ECM). There is increasing evidence indicating that development of invasive tumor phenotype involves altered ionic homeostasis driven by aberrant ion channel expression/function, referred to as ‘oncochannelopathies’ (Prevarskaya *et al*, 2018a). It is now well established that, on the one hand, Ca^2+^ signaling plays a crucial role in cancer development (Monteith *et al*, 2017), while, on the other hand, elevated tissue Na^+^ concentration is a highly specific *in vivo* indicator of malignant lesions in human cancer patients (Haneder *et al*, 2015; Zaric *et al*, 2016). This pinpoints the unraveling of the mechanisms coupling the alteration in Na^+^ homeostasis to intracellular Ca^2+^ events upon cancer progression as one of the central subjects in the field of oncochannelopathies.

Over the years, several studies have revealed some striking similarities in the behavior between neuronal and metastatic cancer cells. Indeed, gene products traditionally known for their role in normal neuronal migration, axonal growth and embryonic development, were also pinpointed in metastasis, where they contribute to the aggressive cancer phenotype (Shah *et al*, 2018; Biankin *et al*, 2012). Interestingly, ion channels responsible for generation of action potential in the electrically excitable cells were also reported in metastatic cells (Djamgoz *et al*, 2019). An intriguing player of neuron-like pathway in metastatic cancer cells seems to be the Na^+^ leak channel, NALCN, first reported in neurons (Lu *et al*, 2007), and whose expression was then found to be altered in some types of cancer (reviewed in (Cochet-Bissuel *et al*, 2014)) with, as yet, unknown consequences for cancer cells behavior. NALCN baseline activity in neurons produces about 10 mV depolarizing shift in their resting membrane potential (V_r_) (Lu *et al*, 2007). However, most importantly, NALCN is part of a channelosome, a multi-protein complex whose members are involved in channel folding, stabilization, cellular localization and activation (Cochet-Bissuel *et al*, 2014) thereby providing great degree of versatility for the regulation of its function not only in neurons, but also in cancer cells.

Another important question is how transmembrane Na^+^ fluxes can promote cancer cell invasiveness. Indeed, it is mainly dysregulated intracellular Ca^2+^ signaling, with the specific spatial and temporal signatures adopted by cancer cells, which is currently acknowledged to drive the progression of certain cancer hallmarks, including invasion and metastasizing (Prevarskaya *et al*, 2018b, 2011). Given the existence of several tightly linked Na^+^/Ca^2+^-dependent molecular pathways that participate in the maintenance of the intracellular Ca^2+^ and Na^+^ homeostasis (Verkhratsky *et al*, 2018), we hypothesized that in metastatic cancer cells, NALCN could provide signaling input encouraging the cell invasion via intracellular Ca^2+^ events.

Thus, in the present work we have undertaken a thorough investigation of the neuronal-type NALCN in prostate cancer cells in the context of its significance for prostate carcinogenesis. Our findings identify NALCN as a major mechanism responsible for Na^+^ influx that governs Ca^2+^ oscillatory signaling (involving cytoplasm, endoplasmic reticulum and mitochondria), ROS production and initiation of invadopodia formation (referred hereafter as invadopodogenesis) in strongly metastatic prostate cancer cells. Importantly, formation of dynamin puncta, reporting invadopodial precursor assembly, occurred at sites of the [Ca^2+^]_c_ wave initiation enriched with NALCN. The NALCN-governed oscillatory Ca^2+^ events facilitate secretion of proteolytic enzymes (matrix metalloproteinases, MMPs) and promote invadopodogenesis via Src kinase activation and actin remodeling (also encouraged by N-WASP – Cdc42 coupling). Modulation of NALCN bioavailability strongly affects *in vivo* prostate cancer progression and bone metastasis formation.

Overall, our study uncovers malignant assignment of NALCN and demonstrates its role as the central element of critical Na^+^/Ca^2+^ signaling axis promoting metastatic progression. Our data suggest that *NALCN* gene must be included in the panel of screened genes for early diagnosis of metastatic prostate cancer.

## Results

### NALCN promotes metastasis-associated cancer cell behavior

Initial immunohistochemical analysis of human prostate tissue arrays detected NALCN upregulation during cancer progression whereas NALCN was not detected in non-cancerous prostate glands (Figure 1a). In clinically localized prostate cancer, NALCN expression was significantly higher within the ISUP group (Gleason score that reflects loss of differentiation) (Table S1). NALCN expression was also significantly higher in pT3 tumors (with extra-prostatic extension) as compared to pT2 tumors (limited to the prostate gland) (Table S1). In human prostate cancer, NALCN immunostaining was observed in 57% of clinically localized hormone-naïve cases, 6% (3/48) of castration-resistant prostate cancer (CRPC) and 62% (13/21) of metastases. The expression of NALCN was strongly correlated with the expression of Src-kinase (Figure 1b). *NALCN* mRNA level was significantly higher in prostate cancer biopsies relative to non-cancerous tissues taken from the same patients (Figure 1c). Similar results were obtained for NALCN expression in other cancers: positive staining in bone metastases from breast (11/19) and colon (5/8) cancers, without any detectable expression in corresponding normal tissues (0/5) (Figure 1d). These data suggest that NALCN expression has clinical relevance and that NALCN may contribute to malignancy in several carcinomas.

**Figure 1:**
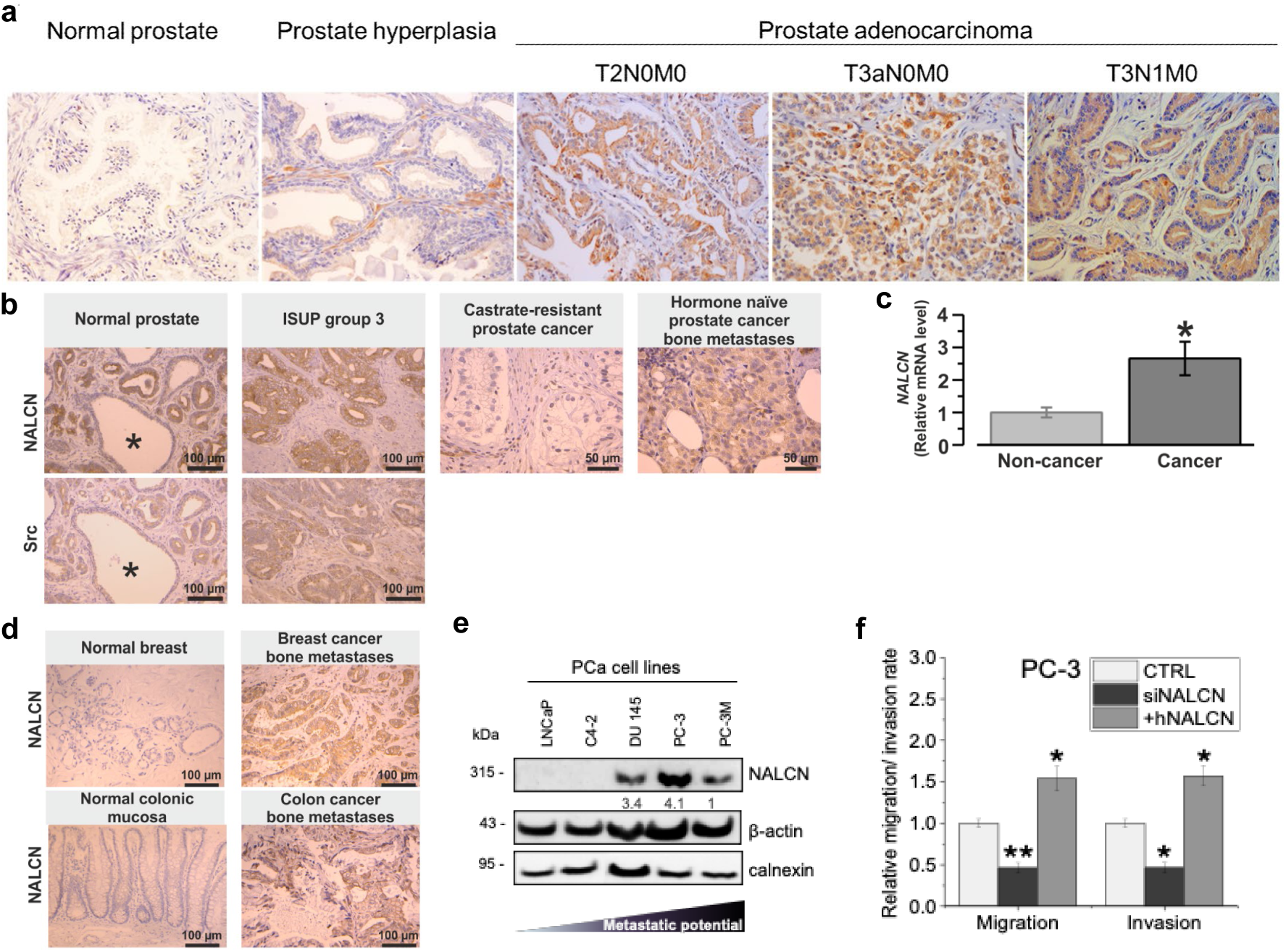
NALCN promotes prostate cancer progression. (a) Immunohistochemical positive staining for NALCN in human prostate adenocarcinoma tissues, but not in normal or hyperplastic prostate tissues (tissue microarray: PR484, US Biomax, Inc). T2N0M0 - tumor invaded submucosa without any regional or distant metastases (stage II); T3aN0M0 – tumor had broken through the capsule of the prostate gland and invaded through muscularis propria (stage III); T3N1M0 – tumor invaded through muscularis propria with 1 to 3 regional lymph node metastases (stage IV). (b) Immunohistochemistry: significant correlation (P<0.0001) between positive staining of NALCN and Src in clinically localized prostate cancer, but not in non-cancerous (*) prostate. Note smooth muscle fiber staining as positive internal control for NALCN. (c) RT-qPCR: NALCN mRNA level in cancerous vs. non-cancerous prostate biopsies (n=28). Data are mean±S.E.M. *P<0.05, two-tailed Mann-Whitney U test. (d) Immunohistochemistry: positive staining of NALCN in bone metastases produced by breast and colon cancers, but not in normal corresponding tissues. (e) NALCN protein level in human prostate cancer cell lines. Data are normalised to β-actin and calnexin and are presented as mean values±SEM of three independent experiments (n=3). (f) Transwell® migration and Matrigel invasion assays after transient NALCN suppression and overexpression. Control cells were transfected for 72 hours whether with siRNA targeting firefly luciferase (CTRL), whereas NALCN depleted cells with siRNA (siNALCN) and NALCN overexpressing cells (+hNALCN). Data are presented as mean±S.E.M of four independent experiments. *P<0.05, two-tailed Mann-Whitney U test.

Consistent with this, screening of various prostate cancer cell lines revealed NALCN and all members of the NALCN channelosome - UNC-79, UNC-80 and NLF-1 - only in highly aggressive cell lines, with little or no expression in weakly metastatic cells (Figures 1e and S1a). Furthermore, NALCN was functionally expressed in the plasma membrane (PM) (Figure S1b), thus providing a pathway for Na^+^ influx, which was not observed in NALCN-negative cell lines (Figures S1c-e). Importantly, NALCN silencing did not affect cellular viability, cell cycle progression, apoptosis (basal or induced) or proliferation (Figure S2), whereas metastasis-associated behaviors (cellular motility, migration and invasion) were significantly dependent on the level of NALCN expression (Figure 1f).

Recruitment of NALCN to invadopodogenesis was studied by cyanine-3 (Cy3)-fluorescent gelatinase assay. NALCN silencing significantly suppressed the number of the invadopodium formation sites and the total area of gelatin degradation (Figures 2a and S2f).

**Figure 2.**
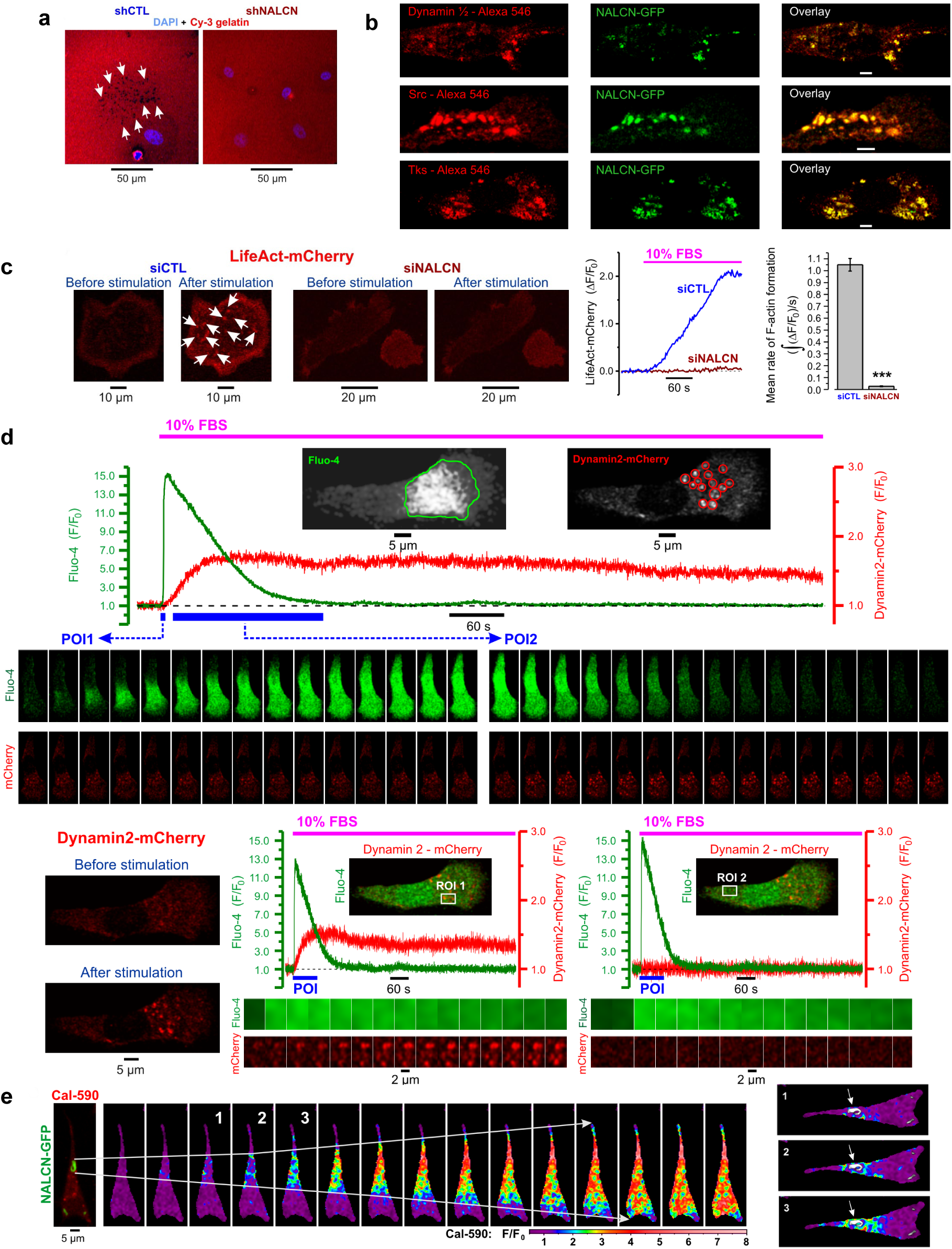
Spatial distribution of NALCN and its signaling input in PC-3 Cells. (a) NALCN knockdown suppresses invadopodium formation reported by Cy3-fluorescent gelatin degradation (arrows). Confocal images of Cy3 (red) and DAPI (blue; nuclei) fluorescence in control (shCTL) and shNALCN 48 hours after plating the cells on gelatin. (b) Co-localisation of NALCN with invadopodia markers: Dynamin (top), Src (middle) and Tks (bottom). (c) NALCN silencing suppresses invadopodial precursor assembly (arrows: F-actin-enriched regions reported by LifeAct-mCherry). Left: near-cell-bottom confocal images captured before and 3 min after stimulation with 10% fetal bovine serum (FBS). Middle: corresponding traces of relative changes in mCherry fluorescence. Right: mean rate of F-actin formation estimated as signal mass () per second in cells transfected with siCTL (n=93) and with siNALCN (n=140). Data: mean±S.E.M. ***P<0.001, two-tailed Student’s t-test. (d) Plot (top) shows temporal profiles of self-normalised fluorescence intensity (F/F0) of fluo-4 (green), averaged within outlined region (inset: left, green outline), and mCherry (red), averaged within 14 circles (inset: right, red outlines). The galleries (bottom) show confocal images of fluo-4 and mCherry fluorescence captured during two periods of interests, POIs (depicted by blue bars on the plot): POI1 (left gallery: every image) and POI2 (right gallery: every 100th image). Bottom left: confocal images of Dynamin2-mCherry fluorescence captured before and 3 min after stimulation with 10% FBS. Middle and right: plots relate the dynamics of [Ca2+]c changes (Fluo-4) to formation of dynamin puncta (Dynamin2-mCherry) at two regions of interests (ROIs) (insets); the galleries (below) show every 40th image (after 90° rotation) during the POIs (plots: blue bars). (e) Top: the overlay of NALCN-GFP and Cal-590 images (left) is related to the gallery of self-normalized rainbow-coded Cal-590 images showing [Ca2+]c response to 10% FBS. Right: 3 rainbow-coded Cal-590 images are superimposed on grey-coded NALCN-GFP image. Arrows: NALCN-enriched structure.

Previously, pro-invasive role of Na^+^ influx through the de novo expressed Na_v_ channels in breast cancer cells was ascribed to facilitated H^+^ extrusion via Na^+^/H^+^ exchanger type 1 (NHE-1), leading to acidification of the pericellular microenvironment and activation of ECM-degrading cysteine cathepsins at invadopodia sites (Gillet *et al*, 2009a; Brisson *et al*, 2013a). Surprisingly, in our experiments, knockdown of Na^+^-conducting NALCN (NALCN-KD) had no effect on either intracellular pH or H^+^ extrusion from the PC-3 cells (Figure S2g). This argued against the possibility that metastasis-promoting significance of NALCN-mediated Na^+^ influx in prostate cancer cells is somehow linked to pH changes.

Immunocytochemical analysis revealed strong co-localization of NALCN with Src kinase, a key signaling component of the invasion pathway, and with invadopodial proteins (cortactin, dynamin, matrix metalloproteinase MT1-MMP and adaptor protein Tks5) within specific morphological structures referred to as ‘invadopodia puncta and rosettes’ (Figures 2b and S2q). At the same time, when PC-3 cells co-expressing NALCN-GFP and Tks5 were placed on gelatin, it was newly formed invadopodia demonstrating the colocalization of NALCN and Tks5 (Figure S3). NALCN targeting to invadopodia was also demonstrated by subcellular fractionation (Figure S2p). Although previous studies have reported that Src kinase is recruited to the NALCN channelosome (Lu *et al*, 2009), no evidence of direct Src kinase-dependence of NALCN expression was presented. We have found that NALCN-KD markedly suppressed both Src kinase expression and activity in metastatic PC-3 cells (Figures S2n and S2o).

Fetal bovine serum (FBS) containing the mixture of various growth factors, or epidermal growth factor alone (EGF) which acts through surface receptor tyrosine kinases (RTKs) or G-protein-coupled receptors (GPCRs) could be used to stimulate pro-invasive behavior of prostate cancer cells cultured under starving conditions. Indeed, both FBS and EGF were reported as potent pro-invasive stimuli (Sun *et al*, 2014). It was previously demonstrated that stimulation of starved melanoma cells with fetal bovine serum (FBS) induces Ca^2+^ oscillations and facilitates invasion. However, the casual link between the two processes remained unknown. We have found that in PC-3 cells, FBS also induced [Ca^2+^]_c_ response, associated with deviation of PI(4,5)P_2_ (phosphatidylinositol 4,5-bisphosphate) from equilibrium, facilitated interaction between N-WASP (neuronal Wiskott–Aldrich syndrome protein) and Cdc42 (cell division cycle 42 protein), and induced invadopodial precursor assembly detectable as enriched F-actin and dynamin puncta (Figure S4). Importantly, this process was strongly suppressed by NALCN-KD (Figure 2c). Consistent with our hypothesis of the existence in invasive cells of a specific Ca^2+^-Na^+^ signaling axis, we have found that formation of dynamin puncta, reporting invadopodial precursor assembly in PC-3 cells, occurred at sites of [Ca^2+^]_c_ wave initiation (Figure 2d) that, in turn, coincided with NALCN-enriched rosettes (Figure 2e).

### NALCN Drives Ca^2+^ Oscillations

We have found that exposure of PC-3 cells to FBS or EGF alone induces an initial [Ca^2+^]_c_ transient followed by [Ca^2+^]_c_ oscillations with reciprocal changes of Ca^2+^ concentration in the endoplasmic reticulum ([Ca^2+^]_ER_) (Figures 3a and 3b). The initial [Ca^2+^]_c_ transient, but not the subsequent [Ca^2+^]_c_ oscillations, persisted in Ca^2+^-free solution supplemented with 0.5 mM EGTA, while both were abolished following Ca^2+^ store depletion with 50 µM cyclopiazonic acid (CPA) even when the cells were bathed in Ca^2+^-containing solution (Figure 3c). We concluded, therefore, that generation of these [Ca^2+^]_c_ oscillations involves both Ca^2+^ entry and Ca^2+^ recycling by the endoplasmic reticulum (ER).

**Figure 3.**
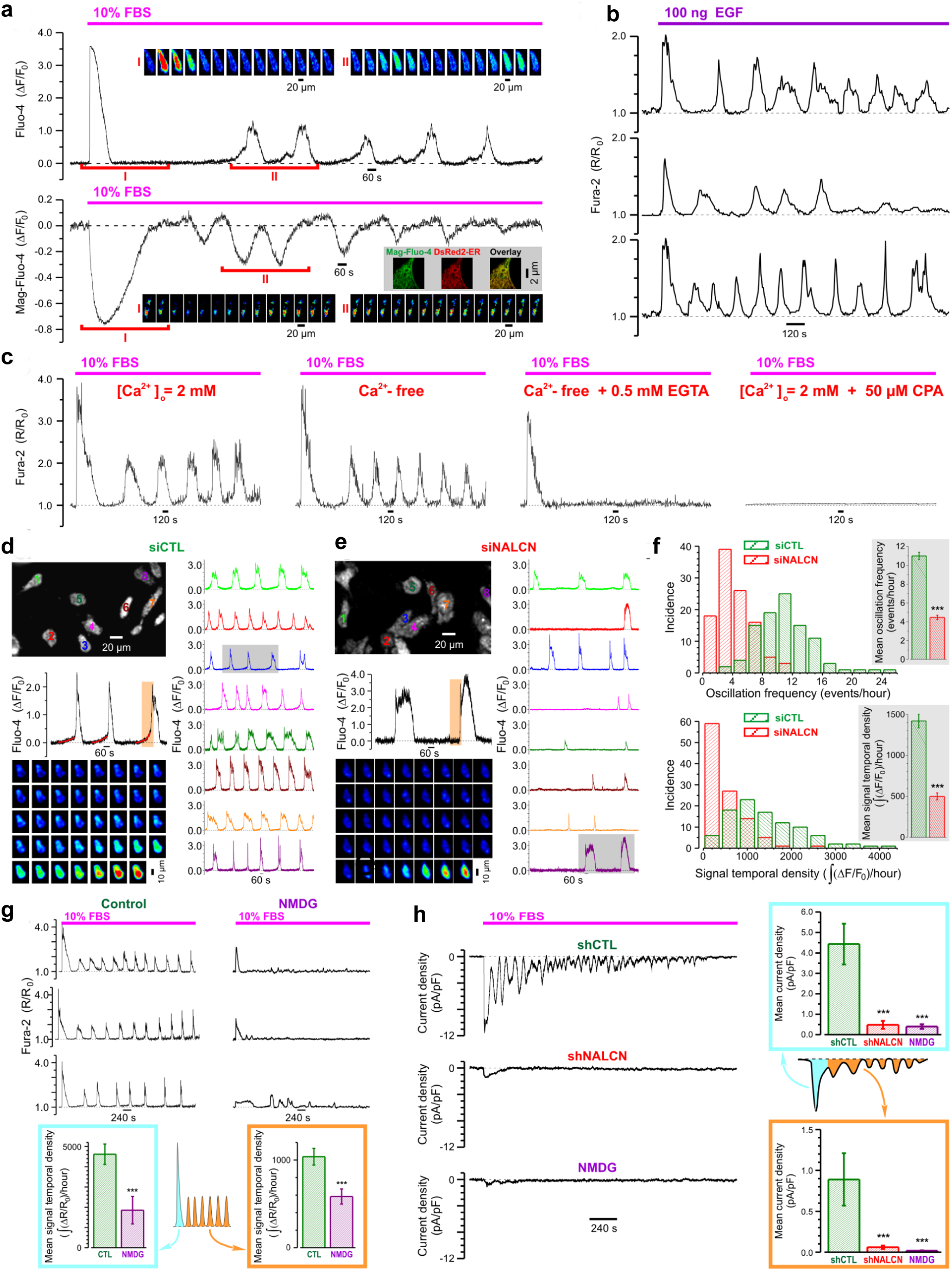
NALCN Regulates Ca2+ Oscillations in PC-3 Cells. (a) FBS-induced transient cytosolic Ca2+ ([Ca2+]c) elevations (Fluo-4, top) are associated with transient decreases of Ca2+ concentration in endoplasmic reticulum ([Ca2+]ER) (Mag-Fluo-4, bottom). The galleries (insets) show every 100th image during the highlighted periods (I and II, respectively). Visualisation of the ER with pDsRed2-ER vector confirms the ER origin of Mag-Fluo-4 signal (grey background inset: confocal images of cell fragment). (b) Sample traces of epithelial growth factor (EGF)-induced [Ca2+]c responses (Fura-2). (c) Contribution of Ca2+ influx and Ca2+ release to FBS-induced [Ca2+]c response (Fura-2) illustrated by sample traces (left to right): control, nominally Ca2+-free or EGTA-supplemented solutions, and Ca2+-containing solution but depleted Ca2+ stores. (d and e) [Ca2+]c oscillations reported by confocal time-series imaging of Fluo-4 fluorescence in FBS-exposed PC-3 cells pre-treated with siCTL (d) or siNALCN (e) for 72 hours. The trace colours (right) correspond to the colour of the cell number (top left). Highlighted (grey background) periods are presented on enlarged plots (left middle) to emphasize that NALCN suppression abolishes “pacemaker events” observed in control (fitted red curves). The galleries (left bottom) show every 3rd image during the highlighted periods (left middle: orange background). (f) The histograms compare the oscillation frequency (top) and signal temporal density (bottom) distributions for siCTL (n=98) and siNALCN (n=107). Insets: corresponding mean±S.E.M. ***P<0.001, two-tailed Student’s t-test. (g) FBS-induced [Ca2+]c response (Fura-2, ΔR/R0) is attenuated following reduction of extracellular Na+ concentration ([Na+]o) to 19.5 mM (substitution with NMDG). (h) FBS-induced inward current (perforated patch, Vh=−80 mV) is suppressed following pre-treatment with shNALCN or reduction of [Na+]o to 13 mM (substitution with NMDG). The bar diagram plots: mean signal temporal densities (g, n=20-23) and mean current densities (h, n=4-6) during initial transient (cyan) and oscillations (orange). Data: mean±S.E.M. ***P<0.001, two-tailed Mann-Whitney U test.

To assess the role of NALCN in the [Ca^2+^]_c_ oscillations, we compared spatiotemporal patterns of the FBS-induced [Ca^2+^]_c_ response in serum-starved control and NALCN-KD PC-3 cells (Figures 3d and 3e). This revealed that the FBS-induced [Ca^2+^]_c_ oscillations (both frequency and signal temporal density) were significantly attenuated in NALCN-KD cells (Figure 3f). Also, the Ca^2+^ “pacemaker phase” (i.e., the phase of slow [Ca^2+^]_c_ rise preceding regenerative response) observed in these oscillations appeared NALCN-dependent as well (Figures 3d and 3e). The [Ca^2+^]_c_ oscillations were attenuated dramatically by reducing extracellular Na^+^ concentration ([Na^+^]_o_) (Figure 3g). Furthermore, application of FBS to PC-3 cells activated an oscillating inward current. This current was dramatically reduced while oscillations were virtually completely abolished following NALCN-KD or [Na^+^]_o_ omission (Figure 3h). These findings were substantiated by a reversal strategy: NALCN overexpression (Figure S5) significantly increased FBS-induced inward current and the [Ca^2+^]_c_ oscillations (Figures S5c-f).

Thus, exposure of serum-starved strongly metastatic PC-3 prostate cancer cells to FBS activates NALCN-mediated Na^+^ influx which appears to be causally linked to generation of [Ca^2+^]_c_ oscillations. Since PC-3 cells exposure to FBS also enhanced Src kinase activity (Figure S6a), one can conclude that FBS-activated NALCN-mediated Na^+^ influx and ensuing oscillatory Ca^2+^ signaling (Figures 3) promote the activity of Src kinase which transduces signals related to cellular proliferation, differentiation, motility and adhesion.

### Coupling between NALCN-mediated Na^+^ influx and [Ca^2+^]_c_ events

Oscillations generally require positive feedback between the contributing elements. We, therefore, assessed (i) modulation of NALCN-mediated Na^+^ influx by extracellular ([Ca^2+^]_o_) and intracellular ([Ca^2+^]_c_) Ca^2+^ concentrations as well as (ii) coupling between Na^+^ influx and rise of [Ca^2+^]_c_.

To unravel the net effect of [Ca^2+^]_c_ elevation on NALCN-mediated Na^+^ influx, we assessed the Na^+^ influx induced by switching [Na^+^]_o_ from 0 to 130 mM (Figure 4a) and by store-operated Ca^2+^ entry (SOCE) (Figure 4c). The two were then compared (Figure 4d, top). We have found that [Na^+^]_o_ rise induced a significant Na^+^ influx (Figure 4a, left) which: (i) was dramatically suppressed by NALCN-KD (see also Figure 4d); (ii) was not modulated by [Ca^2+^]_o_, at least within the range of 0 – 4 mM (Figure 4a, middle; Figure S6c), in contrast to what was reported for neurons (Lu *et al*, 2010); and (iii) was not affected by Ca^2+^ store depletion (Figure 4a right, Figure S6d). Furthermore, [Ca^2+^]_c_ elevation brought about by acute inhibition of sarco/endoplasmic reticulum Ca^2+^-ATPase (SERCA) with thapsigargin (TG) failed to induce Na^+^ influx when the cells were bathed in Ca^2+^-free external solution supplemented with 0.5 mM EGTA, i.e., when SOCE was prevented (Figure 4b). Moreover, we have found (Figure 4c) that: (i) SOCE activation triggers Na^+^ influx even at constant physiological [Na^+^]_o_ in a [Ca^2+^]_o_-dependent manner (see also Figures S6c-f); (ii) NALCN-KD significantly suppresses not only this Na^+^ influx but also SOCE; and (iii) inhibition of the reverse-mode of plasmalemmal Na^+^/Ca^2+^ exchanger (RM-NCX) curtails [Ca^2+^]_c_ transient caused by SOCE at [Na^+^]_o_=130 mM, does not affect it at [Na^+^]_o_=0 mM and attenuates [Ca^2+^]_c_ transient triggered by Na^+^ readmission following SOCE activation. Comparison of the effects of NALCN-KD on Na^+^ influx caused by the [Na^+^]_o_ rise with that triggered by SOCE revealed that SOCE significantly augmented NALCN-mediated Na^+^ influx (Figure 4d, top). In addition, the cell-surface biotinylation assay reported that SOCE increased expression of NALCN protein in the PM (Figure 4d, bottom), similar to FBS (Figure S6g).

**Figure 4.**
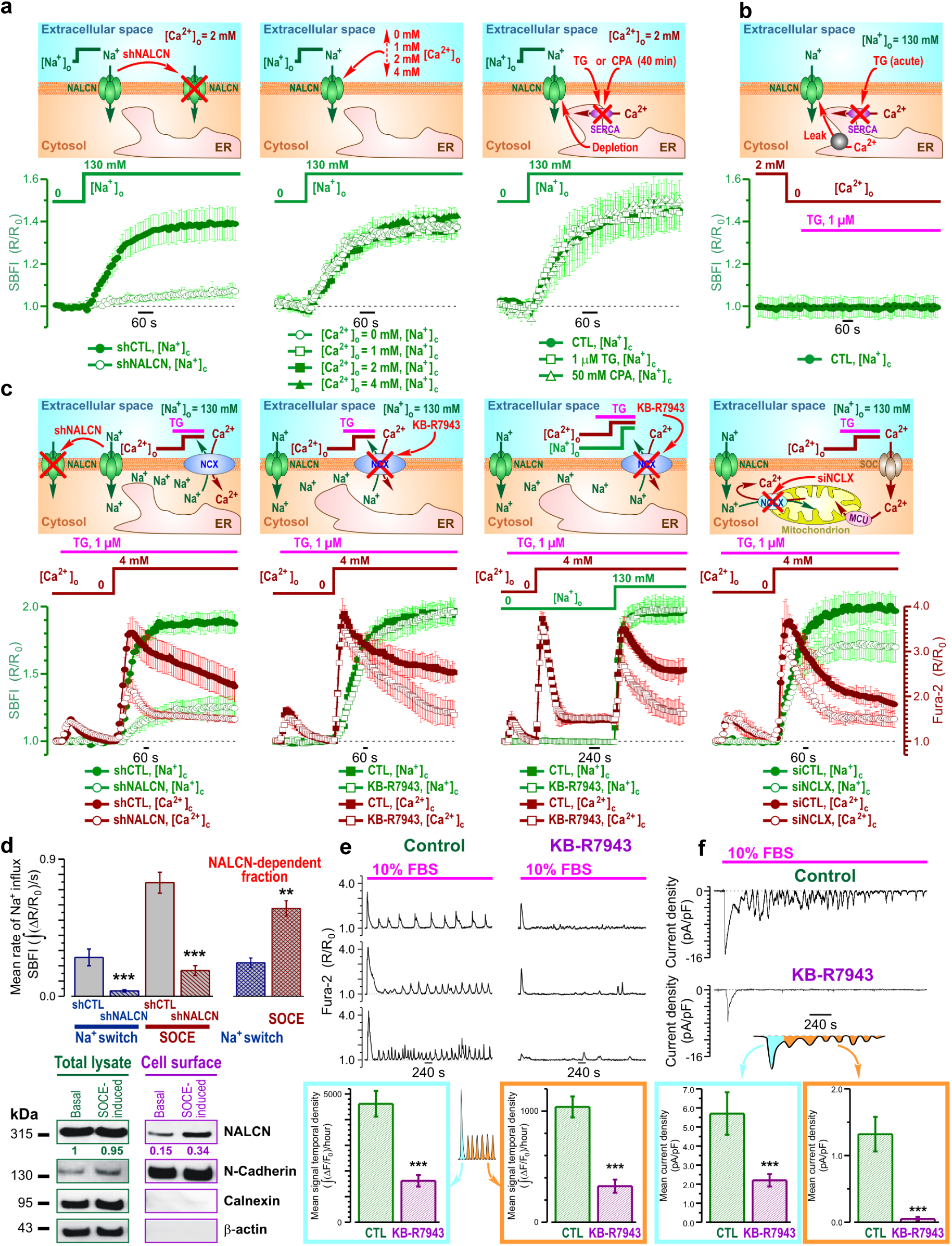
Interplay between SOCE, NALCN, NCX and NCLX in PC-3 Cells. (a and b) Extracellular Na+ concentration ([Na+]o) changes reported by ratiometric Na+ indicator SBFI following (a) [Na+]o switch from 0 to 130 mM and (b) application of 1 µM thapsigargin (TG), the sarco/endoplasmic reticulum (SERCA) inhibitor uncoupling the endoplasmic reticulum (ER) Ca2+ leak from Ca2+ re-uptake into the ER. In (a) the effect of (left to right): NALCN knockdown, extracellular Ca2+ concentration ([Ca2+]o) and Ca2+ store depletion. (c) Store-operated Ca2+ entry (SOCE)-induced changes of cytosolic Na+ concentration ([Na+]c) (SBFI) and cytosolic Ca2+ concentration ([Ca2+]c) (Fura-2); the effect of (left to right): NALCN knockdown, the reverse-mode plasmalemmal Na+/Ca2+ exchanger (RM-NCX) inhibitor KB-R7943 (1 µM) at [Na+]o=130 mM and [Na+]o stepped from 0 to 130 mM following SOCE activation, and knockdown of the mitochondrial Na+/Ca2+ exchanger (siNCLX). In (a-c) cartoons (top) highlight experimental design and plots (bottom) show traces (mean±S.E.M; n=45-501) of SBFI (olive) and Fura-2 (wine) fluorescence with the axes (R/R0) presented in corresponding colour. (d) SOCE addresses NALCN to plasma membrane. Plots (top) compare mean rates of Na+ influx induced by [Na+]o-switch (n=283-360) and SOCE (n=180-216), and their shNALCN-sensitive fractions. Immunoblotting (bottom) compares NALCN expression in total cell lysates and biotinylated fractions before and following SOCE induction. Numbers show (in corresponding colour) mean values (n=3) of NALCN protein levels (band intensity) normalized to N-Cadherin, Calnexin and β-actin. (e and f) KB-R7943 (1 µM) attenuates: (e) [Ca2+]c responses (Fura-2, R/R0) and (f) inward current (perforated patch, Vh=−80 mV) induced by 10% FBS. The bar diagram plots: mean signal temporal densities (e, n=20) and mean current densities (f, n=6-8) during initial transient (cyan) and oscillations (orange). Data are mean±S.E.M. **P<0.01, ***P<0.001, two-tailed Mann-Whitney U test.

The above results indicate that Ca^2+^ influx during SOCE (Figures 4c,d), but not Ca^2+^ release from the ER (Figure 4b) or Ca^2+^ store depletion *per se* (Figure 4a, right), enhances NALCN-mediated Na^+^ influx by promoting translocation of NALCN protein to the PM. In turn, the rise in cytosolic Na^+^ concentration ([Na^+^]_c_) facilitates Ca^2+^ influx either via reverse mode of plasma membrane Na^+^/Ca^2+^ exchanger, RM-NCX (Figures 4c, middle; Figure S6h), and/or by relieving SOCE inactivation. The latter may involve mitochondrial Na^+^/Ca^2+^ exchanger (NCLX), which was shown previously to affect SOCE via modulation of reactive oxygen species (ROS) production (Ben-Kasus Nissim *et al*, 2017). Consistent with this, we have found that NCLX silencing indeed suppressed SOCE (Figures 4c, right, and S6i).

Thus, in view of the obtained results the following sequence of events in PC-3 cells can be proposed: NALCN-mediated Na^+^ influx → NCLX-mediated Ca^2+^ extrusion from mitochondria → decrease of ROS production → relief of SOCE inactivation. At the same time, attenuation of the FBS-induced [Ca^2+^]_c_ events (both the initial transient and oscillations) by the RM-NCX inhibitor KB-R7943 (Figure 4e), suggests that NALCN-mediated Na^+^ influx facilitates RM-NCX-mediated Ca^2+^ entry, while inhibition of FBS-induced inward current by this compound (Figure 4f) indicates that this Ca^2+^ entry encourages NALCN-mediated Na^+^ influx. This positive feedback between NALCN and RM-NCX is most likely the mechanism sustaining the [Ca^2+^]_c_ oscillations. The modulation of the feedback via the ER Ca^2+^ recycling is also tuned by SOCE and RM-NCX (Figure 3a). We also verified that the Na^+^ influx, attributed to the NALCN-mediated current in PC-3 cells, is tetrodotoxin-resistant and gadolinium-sensitive (Figures S6j-o), similar to neuronal NALCN (Lu *et al*, 2007).

### NALCN – NCLX functional coupling recruits mitochondria to regulation of [Ca^2+^]_c_ oscillations

The tight functional coupling between SOCE and NALCN (Figures 4c,d), the dependence of [Ca^2+^]_c_ oscillations on NALCN-mediated Na^+^ influx (Figures 3d-g) and the involvement of NCLX in SOCE regulation (Figure 4c, right) via ROS-mediated oxidation of Ca^2+^ release-activated Ca^2+^ channel protein 1, Orai1 (Ben-Kasus Nissim *et al*, 2017), suggest that, apart from RM-NCX, the NALCN-mediated Na^+^ influx may also regulate [Ca^2+^]_c_ oscillations via NCLX. Indeed, NCLX would exchange mitochondrial Ca^2+^ to Na^+^ delivered via NALCN, and hence oppose Ca^2+^ uptake by the mitochondrial Ca^2+^ uniporter (MCU) and subsequent ROS production. Consistent with this, we observed that the FBS-induced [Ca^2+^]_c_ oscillations were associated with oscillations of (i) Ca^2+^ concentration in mitochondria ([Ca^2+^]_mito_; Figure 5a) and (ii) mitochondrial superoxide (O_2_^•−^) production (Figure 5b). Also, application of H_2_O_2_ (endogenous product of O_2_^•−^) suppressed the [Ca^2+^]_c_ oscillations (Figure 5c).

**Figure 5.**
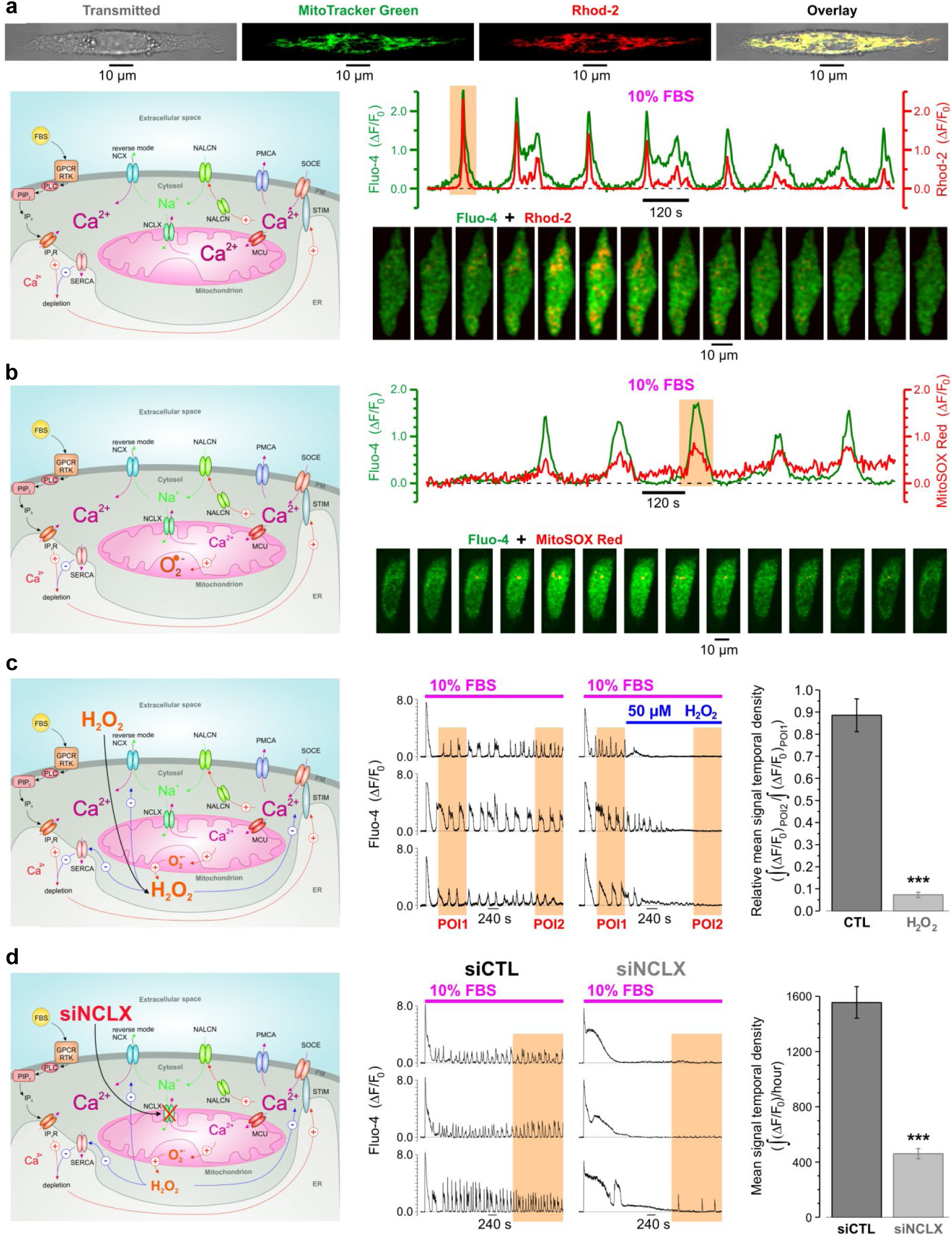
NALCN and Recruitment of Mitochondria to [Ca2+]c Oscillations in PC-3 Cells. Cartoons (left) highlight experimental design and assessed signaling pathways. Note that Na+ delivered to the cytosol by NALCN, on the one hand, activates reverse mode (RM-) of NCX, while on the other hand, is exchanged by NCLX to Ca2+ thus restricting elevation of [Ca2+] in mitochondria ([Ca2+]mito) and, hence, production of superoxide (O2-) and H2O2, which is known to inhibit SERCA, RM-NCX and SOCE elements. (a) Top: visualisation of mitochondria with MitoTracker Green confirms mitochondrial origin of Rhod-2 signal. Middle: Fetal bovine serum (FBS)-induced [Ca2+]c oscillations (Fluo-4) are associated with [Ca2+]mito oscillations (Rhod-2). Bottom: every 2nd image from the highlighted (middle) period. (b) Top: FBS-induced [Ca2+]c oscillations (Fluo-4) are associated with oscillations in O2-production by mitochondria (MitoSOX Red). Bottom: every 3rd image from the highlighted (middle) period. (c) O2-metabolite H2O2 suppresses FBS-induced [Ca2+]c oscillations (Fluo-4). Left: representative traces. Right: mean signal (Fluo-4) temporal density during POI2 was normalized to that during POI1 in control (n=58) and in 50 µM of H2O2 (n=74), and compared. (d) Suppression of NCLX with siRNA (siNCLX) attenuates [Ca2+]c oscillations (Fluo-4). Left: representative traces. Right: mean signal (Fluo-4) temporal densities during highlighted periods were compared in control (n=18) and following 48 h pre-treatment with siNCLX (n=17). Data are mean±S.E.M. ***P<0.001, two-tailed Mann-Whitney U test.

In support of the above hypothesis, we have also found that NCLX silencing (siNCLX, S2T) dramatically attenuated the FBS-induced [Ca^2+^]_c_ oscillations (Figure 5d). Interestingly, the initial [Ca^2+^]_c_ transient caused by the FBS application had significantly longer duration in cells pre-treated with siNCLX, likely due to suppression of SERCA by enhanced ROS production in mitochondria lacking NCLX (Qin *et al*, 2014). Thus, interplay between Ca^2+^ and Na^+^ signals not only would affect mitochondrial Ca^2+^ shuttling and mitochondrial redox status (Ben-Kasus Nissim *et al*, 2017), but also serve to maintain persistent [Ca^2+^]_c_ oscillations. In turn, the latter are necessary for Src activation (Sun *et al*, 2014).

### NALCN regulates secretion of ECM-degrading enzymes in Ca^2+^-dependent manner

In cancer cells, secretion of ECM-degrading enzymes is the principal mechanism facilitating extracellular proteolysis and invasion (Friedl & Wolf, 2008). To assess a role of NALCN in regulation of secretion in aggressive prostate cancer cells we have analyzed the effect of NALCN bioavailability on the rate of FBS-induced secretion visualized by dynamic 3D confocal imaging (Figure 6a) of FM1-43 fluorescent reporter pre-accumulated in the secretory vesicles of starved PC-3 cells. We have found that presence of NALCN proved to be critical for controlling both the rate of vesicle secretion by PC-3 cells (Figures 6b,c) and the activity of ECM-degrading enzymes (Figure 6d). Furthermore, by comparison of the effect of Ca^2+^ chelators with different dynamic properties on the rate of this secretion, we have demonstrated that this process is not only Ca^2+^-dependent, but is, in fact, regulated by [Ca^2+^]_c_ in microdomains, since both fast and slow Ca^2+^ chelators decreased the rate of secretion to different extents (Figures 6e,f).

**Figure 6.**
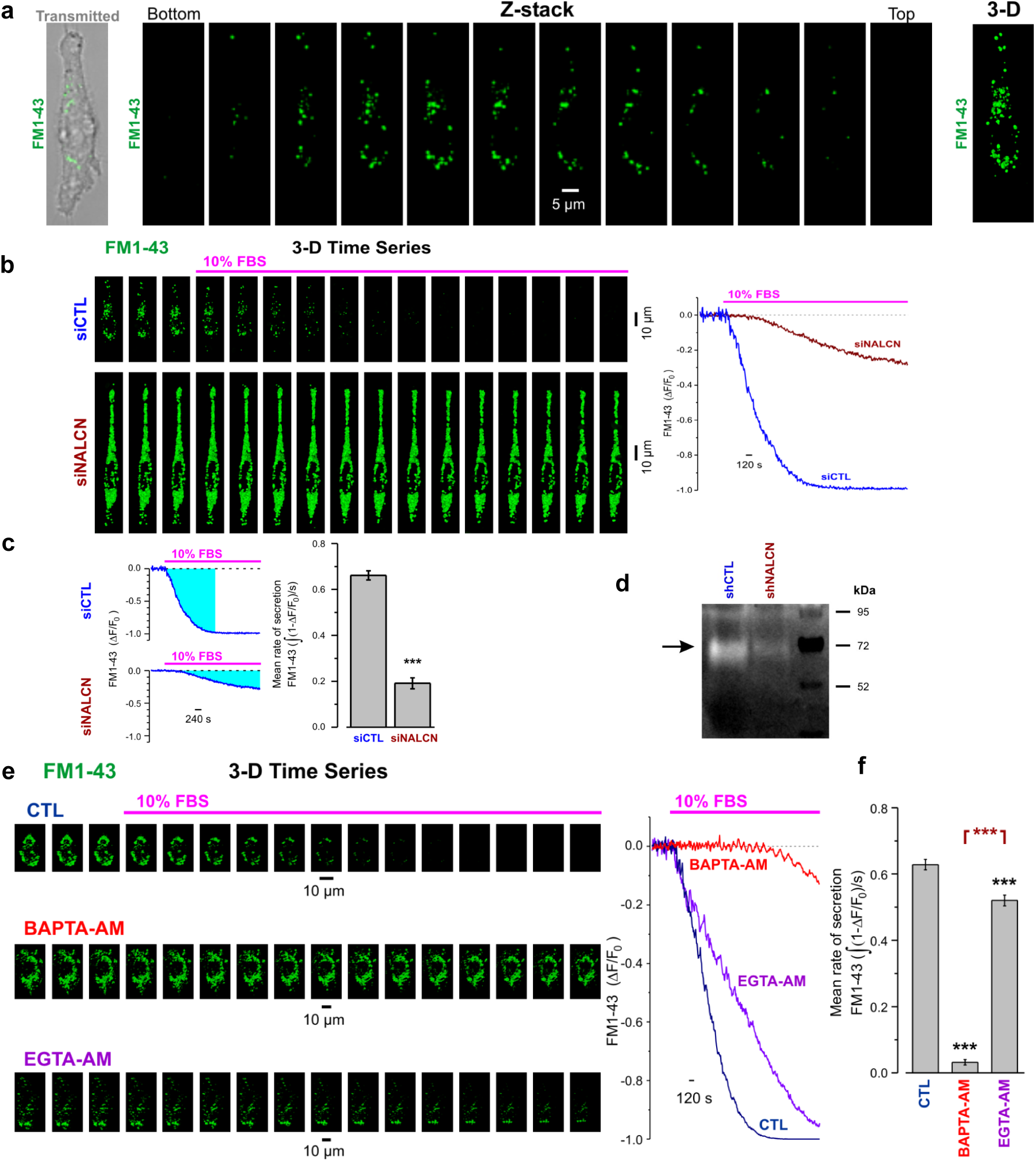
NALCN Regulates Vesicular Secretion of ECM-Degrading Enzymes in Ca2+-Dependent Manner. (a) 3-D distribution of FM1-43-labelled secretory vesicles in PC-3 cell. Left: overlay of transmitted and x-y confocal images of FM1-43 fluorescence. Middle: confocal z-sections. Right: corresponding 3-D image. (b) Left: the galleries of sequential 3-D images (every 25th image) compare FBS-induced degradation of FM1-43-labelled secretory vesicles in PC-3 cells pre-treated with siCTL and siNALCN. Right: corresponding traces of relative changes in total z-stack fluorescence. (c) Plot compares mean rates of secretion, calculated as signal mass (cyan, left) per second, in control (siCTL: n=35) and following NALCN suppression (siNALCN: n=40). (d) Zymography compares gelatinase activity (72 kDa) in shCTL and shNALCN. (e) The same as (b) but for untreated (CTL) cells and those loaded with slow (EGTA-AM) and fast (BAPTA-AM) Ca2+ chelators. (f) The same as (c) but compares the mean rates of secretion in CTL (n=45) and after treatment with EGTA-AM (n=35) or BAPTA-AM (n=62). Data are mean±S.E.M. ***P<0.001, two-tailed Student’s t-test.

### NALCN promotes prostate cancer aggressiveness *In Vivo*

Finally, we have analyzed the role of NALCN in cancer progression *in vivo* by using several mice models. Since phosphatase and tensin homologue (PTEN) is one of the most commonly deleted/mutated tumor suppressor genes in human prostate cancer, we utilized the PTEN-knockout mouse with conditional gene inactivation (floxed allele; L2) as our first *in vivo* model (Parisotto *et al*, 2018). Following prolonged treatment with tamoxifen, PTEN^−/−^ mice produced prostate adenocarcinoma, whereas PTEN^−/−^p53^−/−^ developed more aggressive and invasive phenotype of prostate cancer. *Ex vivo* analyses performed on these prostate tumors revealed that NALCN was markedly upregulated in PTEN^−/−^p53^−/−^ tumors (i.e., during progression of invasive adenocarcinoma), compared with the PTEN^−/−^ tumors (i.e., during initial tumorigenesis) (Figure 7a).

**Figure 7.**
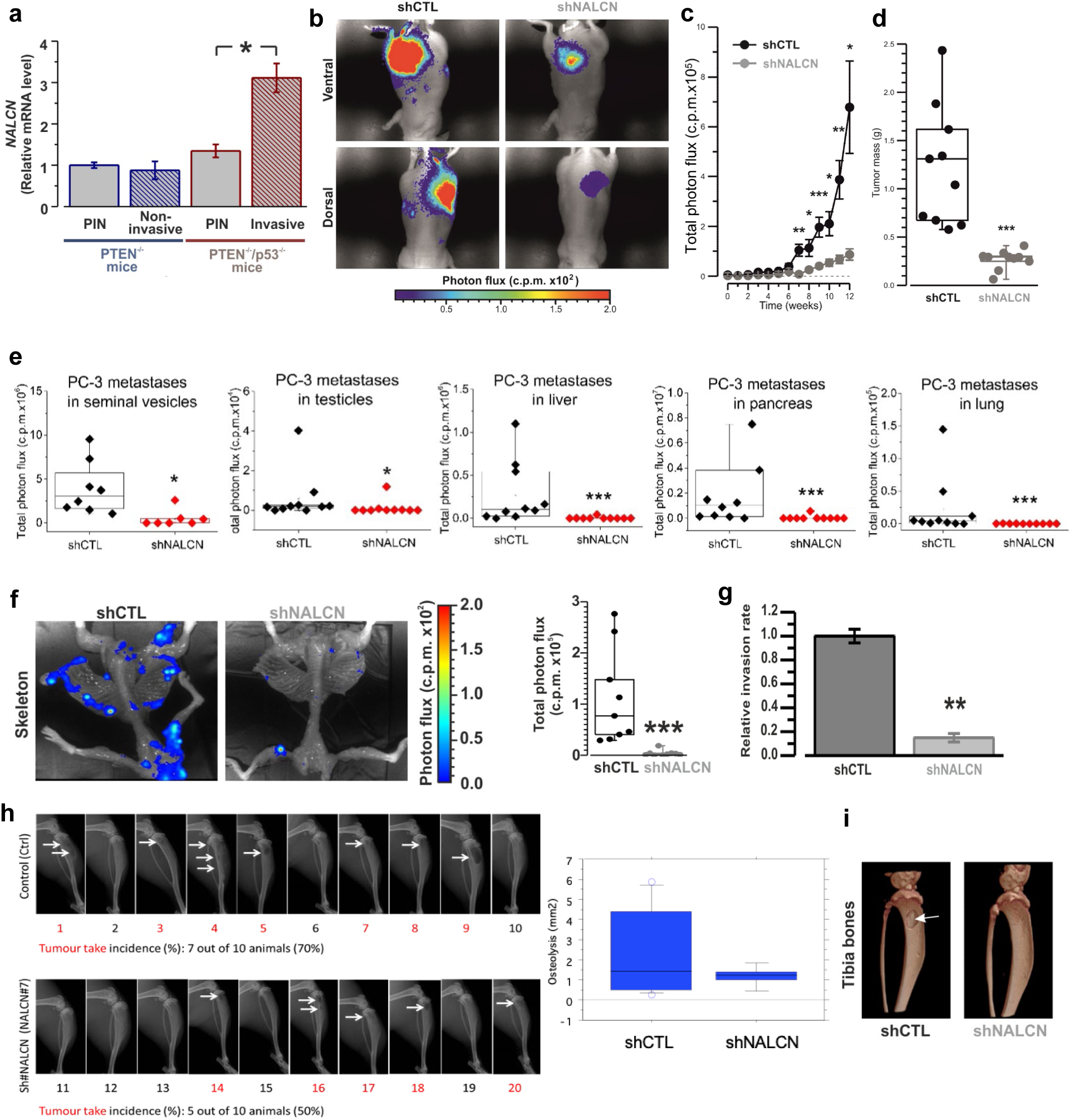
In vivo and ex vivo analysis of NALCN tumorigenesis and metastatic spread. (a) NALCN mRNA level was not different between prostatic intraepithelial neoplasia (PIN) and non-invasive prostate adenocarcinoma of PTEN(i)pe−/− mice. In contrast, invasive adenocarcinoma of PTEN/p53(i)pe−/− mice exhibited significant upregulation of NALCN mRNA when compared with corresponding PIN. Data from PTEN(i)pe−/− mice were normalized to control PTEN(i)pe+/+ mice, whereas data from PTEN/p53(i)pe−/− mice - to control PTEN/p53(i)pe+/+ mice. RT-qPCR data are normalized to GAPDH and TBP and presented as mean±SEM (n=5). *P<0.05, two-tailed Mann-Whitney U test (b-c) Representative images (b) and analysis (c) of bioluminescence imaging (BLI) on mice after orthotopic injections of control (PC-3 Luc-shCTL) or NALCN knockdown cells (PC-3 Luc-shNALCN) revealed tumor growth suppression by NALCN knockdown (n=10). *P<0.05, **P<0.05, ***P<0.001, two-tailed Mann-Whitney U test. Counts per minute (c.p.m.). (d) Masses of tumors dissected from mice used in (c) 12 weeks after the injection: box-plots with raw data (n=10). ***P<0.001, two-tailed Mann-Whitney U test. (e) BLIs in mice organs. Counts per minute (c.p.m.). Box-plots with raw data from each mouse are shown. n.s. – non-significant, *P<0.05, ***P<0.001, two-tailed Mann-Whitney U test. (f) Ex vivo BLIs of skeleton metastases in mice 12 weeks after orthotopic injections of control (PC-3-Luc-shCTL) or NALCN depleted cells (PC-3-Luc-shNALCN). Right: box-plots with raw data of the BLI bone metastasis assay (n=10). ***P<0.001, two-tailed Mann-Whitney U test. (g) NALCN knockdown in PC-3 cells suppresses Transwell® (chemoattractant: conditioned media from osteoblasts) invasion from four independent experiments (n=4). Data are mean±S.E.M. **P<0.01, two-tailed Mann-Whitney U test. (h) X-ray images: osteolysis (arrow) 31 days after intra-tibial injection of control (PC-3 Luc-shCTL) or NALCN depleted cells (PC-3 Luc-shNALCN) with corresponding area (right) of osteolysis (mm2). Data: mean±S.E.M. (i) X-ray 3D images: osteolysis (arrow) 31 days after intra-tibial injection of PC-3-Luc-shCTL or PC-3-Luc-shNALCN cells.

For studying the progression of prostate tumors *in vivo*, we developed xenografts of PC-3 cells with stable luciferase activity. Therefore, our second *in vivo* model was presented by the nude mice that were injected orthotopically with these cells and followed for 12 weeks, allowing primary tumor growth and metastases development. We found that using NALCN-KD PC-3 cells resulted in significantly suppressed metastasis and primary tumor growth *in vivo* (Figures 7b-d). This was confirmed further by *ex vivo* analyses (Figure 7e). It is well known that prostate cancer frequently metastasizes to bone (in fact, the PC-3 cell line was originally derived from the bone metastases of a prostate cancer patient). We observed that injection of NALCN-KD PC-3 cells caused significantly less skeletal metastases compared to the wild-type cells (Figure 7f). NALCN-KD also induced a reduction of metastasis to other organs, albeit to a lesser extent (Figure 7e).

The above observations supported our hypothesis that NALCN expression determines metastatic potential of PC-3 cells, specifically to the bone. Indeed, Transwell^®^ invasion assay utilizing osteoblast conditioned media as a chemoattractant showed significant reduction of the invasiveness of PC-3 cells with NALCN-KD (Figure 7g). Furthermore, the direct injection of PC-3 cells into tibia of nude mice (third *in vivo* model) revealed that the cells with NALCN-KD produced significantly less bone tissue destruction, compared with control wild-type cells (Figures 7h,i and Figure S7). In contrast, assessment of the effect of intracardiac injection of permanently NALCN-overexpressing PC-3 cells (fourth *in vivo* model) revealed that if in control group (PC-3 Luc) only 18% of mice (2/11) developed one unique osteolytic lesion (both were located in the fibula) then with the cells stably overexpressing NALCN (+hNALCN) as high as 64% of mice (7/11) developed multiple osteolytic lesions of larger volumes (Figures S7c,d) in different bone locations (fibula, tibia, hip, humeral, skull and jaw bones).

Overall, the results from all four *in vivo* mice models were consistent, indicating that the level of NALCN expression directly correlated with the metastatic impact of PC-3 derived tumor xenografts.

## Discussion

Present study was undertaken to assess pro-invasive potential of neuronal-type Na^+^ leak channel, NALCN, in prostate carcinogenesis using complex multidisciplinary approach involving: (i) prostate cancer-derived cell lines for elucidating mechanistic aspects of NALCN involvement in cellular signaling promoting invasive behaviors, (ii) *in vitro* animal models for determining metastatic impact of NALCN on tumor xenografts and (iii) primary human prostate cancer biopsy tissues for correlating NALCN expression with clinical tumor scoring and staging. In this study we report on four major findings of the utmost importance for understanding the genesis of the prostate cancer invasiveness: 1) NALCN is overexpressed in highly metastatic prostate cancer cell lines and primary human prostate cancer metastasis; 2) NALCN-mediated Na^+^ entry in strongly metastatic prostate cancer cell lines governs intracellular Ca^2+^ oscillations and promotes invadopodia formation and secretion of ECM-degrading proteolytic enzymes; 3) activation of Na^+^ entry via NALCN engages signaling pathway involving increased mitochondrial Ca^2+^ extrusion via NCLX, decreased mitochondrial ROS production, relief of SOCE inactivation by ROS, augmentation of plasma membrane SOCE and enhancement of Src kinase activity; 4) in the *in vivo* mice models NALCN expression determines metastatic potential of xenograft tumors and formation of bone metastasis.

NALCN channel was first discovered in neurons where its activity provides for constitutive influx of Na^+^ which depolarizes V_r_ by about 10 mV compared to K^+^ equilibrium potential (Lu *et al*, 2007). Subsequently, *NALCN* gene expression was also detected in several types of human cancers (reviewed in (Cochet-Bissuel *et al*, 2014)). However, its functional significance in carcinogenesis remained unknown. Expression of neuronal-type channel, NALCN, in non-excitable cancer cells, in general, is of no surprise in view of the fact that cancer cells’ channelosome is known to undergo essential modification characterized by acquisition of ion channel types inherent to excitability (Prevarskaya *et al*, 2018b). Thus, it is much more important to establish what cancer hallmark(s) newly expressed channel(s) would promote and what intracellular signaling would be engaged.

Among the ion channels of cancer cells inherent to excitability, the voltage-gated, Na^+^-conducting ones of Na_v_1 family, are specifically linked to cancer cells invasion and metastasis (House *et al*, 2010a; Roger *et al*, 2007). It is believed that small steady-state “window current” through these channels is the major contributor to the cancer cells’ depolarized V_r_ and increased [Na^+^]_c_ (Campbell *et al*, 2013; Gillet *et al*, 2009b; Roger *et al*, 2003) with both being the factors suggested to play a role in the mechanism of pro-invasive Na_v_1 channels action. For instance, depolarization of the V_r_ due to Na_v_1.5 activity in colon cancer cells was implicated in the transcriptional induction of invasion-related genes through protein kinase A (PKA), Rap1B, MEK, ERK1/2 signaling pathway (House *et al*, 2015, 2010b), whereas in breast cells the same was linked to modulation of F-actin polymerisation and invadopodia formation (Brisson *et al*, 2013b). Furthermore, in breast cancer cells, Na_v_1.5-mediated Na^+^ entry was shown to underlie intra- and extracellular pH changes via Na^+^/H^+^ exchanger type 1 (NHE1) which in turn favored higher activity of secreted ECM-digesting proteases, cathepsins, facilitating breast cancer cells invasiveness (Brisson *et al*, 2013b).

Taking the above into account, with the discovery of NALCN in cancer cells, an essential part of depolarized V_r_ and enhanced Na^+^ influx in these cells could be ascribed by one to the function of this channel. However, our results show that in highly metastatic prostate cancer cells, coupling NALCN-mediated Na^+^ entry to the intracellular oscillatory-type Ca^2+^ signaling is responsible for the promotion of invasive behavior via invadopodia formation and secretion of ECM-degrading proteolytic enzymes.

A number of Na^+^-permeable channels were suggested to influence [Ca^2+^]_c_ events in different cell models (Verkhratsky *et al*, 2018). In cancer cells, functional coupling of Na_v_ channels activity to intracellular Ca^2+^ signaling was predicted in human colon cancer (House *et al*, 2010a). Later, it was shown that spontaneous [Ca^2+^]_c_ oscillations in strongly metastatic human prostate and breast cancer cells were inhibited by Na_v_ channel blockers (Rizaner *et al*, 2016). However, the molecular mechanisms coupling Na^+^ influx to [Ca^2+^]_c_ oscillations as well as their role in metastatic cell behavior remain largely unknown. The importance of Ca^2+^ signaling in invasion and invadosome function is currently emerging. Yet, the identification of the specific roles of ion channels at different steps of complex adhesive and degradative functions of invadosomes remains challenging. For example, it has been recently shown that TRPV4 calcium channel colocalizes with β1-integrins at the invadosome periphery and regulates the coupling of acto-adhesive and degradative functions(Vellino *et al*, 2021).

In prostate cancer cells we have unraveled a novel, previously not anticipated pathway composed of several feedback elements through which NALCN-mediated Na^+^ influx governs the oscillatory pattern of intracellular Ca^2+^ signaling, which then translates into invasive behavior (Figure S8). Importantly, we have found that this pathway involves mitochondrial Ca^2+^ handling, NCLX and ROS production which were recently referred as the important determinants of cancer growth and metastasis (Verkhratsky *et al*, 2018; Sun *et al*, 2014). This pathway (Figure S8), that could be initiated by the EGF-mediated stimulation of RTK/GPCR, involves initial Ca^2+^ store depletion via derivation of Ca^2+^-mobilizing second messenger, IP_3_, with ensuing activation of transmembrane SOCE. The resulting elevation of [Ca^2+^]_c_ facilitates NALCN translocation to PM channelosome to enable NALCN-mediated Na^+^ entry and increase of [Na^+^]_c_ which then promotes additional Ca^2+^ influx via RM-NCX. Significant rise of [Ca^2+^]_c_ triggers Ca^2+^ uptake back into the ER via SERCA pump and MCU-mediated Ca^2+^ uptake into mitochondria. The latter facilitates production of ROS, which is known to inhibit SERCA (Patel & Brackenbury, 2015), RM-NCX (Liu & O’Rourke, 2013) and SOCE elements (Bogeski *et al*, 2012), and whose action is opposed by NCLX exchanging NALCN-delivered Na^+^ to mitochondrial Ca^2+^. This positive and/or negative feedback underlies generation of [Ca^2+^]_c_ oscillations maintaining the activity of Src, which is an essential component of NALCN channelosome. Active Src, in turn, phosphorylates downstream proteins (cortactin, dynamin and Tks5) recruiting them to actin polymerization regions and giving rise to invadopodia. RTK/GPCR activation is also linked to activation of the Rho family GTPase, Cdc42, leading to its binding with N-WASP and subsequent actin nucleation (Beaty & Condeelis, 2014). In addition, invadopodia maturation is facilitated by Ca^2+^-dependent secretion of ECM-degrading MMPs.

The validity of this NALCN-dependent pathway was confirmed in the *in vivo* experimentation on mice xenograft tumors wherein xenografts produced by NALCN-KD cells were characterized by significant downsizing of skeletal metastases, reduction in bone tissue destruction and diminution of metastasis to other organs.

## Conclusion

In this study, we identified for the first time that the expression of NALCN in highly invasive metastatic human prostate carcinoma. We have found that NALCN is functional and importantly promotes invasiveness and metastatic potential of prostate cancer cells by controlling the Na^+^/Ca^2+^ signature. Specifically, we demonstrated a novel NALCN-mediated mechanism of cancer cell aggressiveness whereby long-term intracellular Ca^2+^ oscillations are the central players. These oscillations are controlled mainly by SOCE, NCX, NCLX, SERCA and ROS, and are necessary for Src kinase activation, invadopodia formation and Ca^2+^-dependent secretion of ECM degrading enzymes. (via PIP2-WASP crosstalk). Thus, by inducing long-lasting Ca^2+^ oscillations following by actine remodeling and MMP secretion, NALCN not only initiates the invasion process, but also provides persistent signaling input augmenting invasiveness of aggressive cancer cells over long period of time.

Furthermore, our results obtained in several mice models suggest that NALCN expression promotes the metastatic impact of prostate cancer *in vivo*.

In overall conclusion, our study identifies a novel ionic checkpoint during the metastatic transformation and pinpoint NALCN as a key player governing this process in prostate cancer.

## Materials and methods

### 1. Cell culture

LNCaP, DU 145 and PC-3 were from the American Type Culture Collection (ATCC^®^). C4-2 and PC-3M were kindly provided by Dr Florence Cabon (Cancer Research Centre of Toulouse, France) and by Dr Scott Fraser and Prof. Mustafa Djamgoz (Imperial College London, UK), respectively. In accordance with their origin / tumorigenic potential these cancer cell lines are classified as: LNCaP – lymph node / weakly metastatic (Horoszewicz *et al*, 1980); C4-2 – chimeric tumor induced by co-inoculating castrated mouse with LNCaP-derived subline and bone fibroblasts / weakly metastatic (Wu *et al*, 1994); DU 145 – brain / moderately metastatic (Stone *et al*, 1978); PC-3 –bone / highly metastatic (Kaighn *et al*, 1979); and PC-3M – liver metastases induced by inoculating mouse with PC-3 / highly metastatic (Kozlowski *et al*, 1984). Cells were cultured at 37°C in RPMI 1640 medium (Gibco™, Thermo Fischer Scientific) supplemented with 10% fetal bovine serum (Gibco™) and 2 mM L-Glutamine (Gibco™). Mouse-derived osteoblast precursor cell line MC3T3 E1 was cultured in MEMα medium (Gibco™) containing 10% fetal bovine serum and 1% Penicillin-Streptomycin (Gibco™).

### 2. Cell transfection

Cells were transfected with 2 µg of corresponding construct or 50 nM of siRNA and 0.2 µg of pmax GFP using either Nucleofector (Amaxa), or X-tremeGene HP DNA Transfection Reagent (Roche), or HiPerFect Transfection Reagent (QIAGEN).

### 3. Cloning procedures

The human NALCN cDNA was previously described (Swayne *et al*, 2009) and the mCherry cDNA was obtained from the pAAV-EF1a-mCherry plasmid (Addgene #114470). The human NALCN cDNA and the mCherry cDNAs were subcloned in the pLV-EF1a-IRES-Blast (Addgene #85133) using standard molecular biology techniques.

### 4. Lentiviral transduction

Lentiviruses were made at the Vectorology facility of Montpellier (https://www.biocampus.cnrs.fr/index.php/en/). Briefly, replication-deficient lentiviruses were produced and titrated as described by co-transfection of the resulting constructs in HEK-293T cells with the HIV-1 packaging plasmid psPAX2 (Addgene #12260) and the plasmid pMD2.G (Addgene #12259) that encodes the vesicular stomatitis virus glycoprotein envelope. Viruses were harvested and then concentrated. Titers were calculated as described (Naldini *et al*, 1996). Once PC-3 cells that were seeded in 35 mm petri dishes reached the confluence of 80-90 % they were transduced with 120.4 µg p24 of the lentiviral suspension.

### 5. Establishment of stable cell lines

48 hours after transfection/ transduction PC-3 cells were cultured with antibiotics (700 µg/ml G418; 2 µg/ml puromycin; 5 µg/ml blasticidin S). The establishment of stable cell lines was confirmed by RT-qPCR and immunoblotting analysis. In order to verify the luciferase activity 150 µg/ml of D-luciferin was used.

### 6. Chemicals

Chemicals were purchased from Invitrogen Life Technologies and Sigma-Aldrich Chemicals unless stated otherwise.

### 7. RNA purification and reverse transcription polymerase chain reaction

Organ tissues and cells were preserved in RNAlater™ (Sigma). Cellular RNA was purified with the NucleoSpin RNA Plus Kit (Macherey-Nagel). Organ tissues were homogenized with Precellys^®^ Tissue Homogenizer and purified with TRIzol™ Reagent (Invitrogen). The cDNA from was obtained with the RNA reverse transcription (Applied Biosystems) following the DNase treatment (Ambion). Conventional RT-PCR was performed by using AmpliTaq Gold^®^ (Applied Biosystems) on C1000 Touch™ Thermal Cycler (Bio-Rad), whereas quantitative (RT-qPCR) - with SsoFast™ EvaGreen^®^ (Bio-Rad) on CFX96 Touch™ Real-Time PCR Detection System.

### 8. Immunoblotting

The cells were lysed in RIPA buffer containing protease and phosphatase inhibitors, sonicated for 10 seconds and centrifuged at 13000 g for 10 minutes in order to eliminate nuclei. The concentration of protein lysate was measured with Pierce^®^ BCA Protein Assay (Thermo Fisher Scientific). 1xSample Reducing Agent and 1xLDS Sample Buffer (Bolt™, Thermo Fisher Scientific) were added to 50 μg of protein lysate and heated at 95^0^C for 5 minutes. Proteins were resolved due to 3-8% Tris-acetate SDS-PAGE and transferred onto methanol-activated PVDF membrane with Pierce G2 Fast Blotter (Thermo Fisher Scientific). Afterwards, the blots were blocked with 5% milk TBS-T buffer (15 mM Tris-HCl, 140 mM NaCl, 0.05% Tween20^®^, pH 7.4) for 1 hour, and then incubated with antibodies. For HRP-conjugated secondary antibodies blots were developed using SuperSignal™ West Dura Extended Duration Substrate (Thermo Fisher Scientific) Images were captured with Amersham Imager 600 (GE Healthcare Life Sciences).

### 9. Cell cycle analysis

Cells were fixed with ice-cold 70 % methanol for 30 minutes, resuspended in 100 µl PBS with 500 µg/ml Ribonuclease A and stained with 50 µg/ml of propidium iodide for 1 hour at 4^0^C. At least 10,000 events per condition were processed by CyAn™ ADP Analyser FACScan flow cytometer (Becton-Dickinson).

### 10. Cellular viability

Cellular viability was evaluated with Trypan Blue Exclusion assay (Thermo Fisher Scientific).

### 11. Apoptosis assay

The level of apoptosis was estimated from the number of apoptotic nuclei revealed with Hoechst staining.

### 12. Proliferation assay

Cells were plated into 96-well plate (10^3^ cells/well) and left overnight at standard growth conditions. At each desired time point, cells were fixed with 50 % Trichloroacetic acid at 4^0^C for 1 hour. After removing fixative solution, cells were stained with 0.4 % Sulforhodamine B (Santa Cruz) in 1 % acetic acid. The dye taken up by cells was then dissolved in 10 mM Tris-base, pH 10.5. The absorbance was measured at 560 nm with TriStar^2^ Multimodal Reader LB942 (Berthold Technologies).

### 13. Time-lapse video microscopy

Cells were seeded at low density, stained with Hoechst and kept at 37°C under 5% CO_2_ in an incubator chamber for time-lapse video recording. Cell movements were monitored with an inverted microscope (Biostation; Nikon) using a 20× objective lens. To enable tracking over long distance with high resolution, the tiling mode (4×4 or 8×3 tiles) of the microscope was used. The entire images were then reconstructed using FiJi Grid/Collection Stitching Plugin (Schindelin *et al*, 2012).

### 14. Cellular motility

Images were acquired every 30 minutes during 48 hours. Image stacks were then analyzed using Trackmate ImageJ plugin (Tinevez *et al*, 2017). For each condition, segmentation was performed on Hoechst stained nucleus. The same optimized tracking parameters were used for each image series. Cells were then automatically tracked using the “simple lap tracker” algorithm and, tracks coordinates and individual track mean velocities were measured for each cell. Only cells tracked over a period of more than 5 hours were taken into account and at least 1000 cells tracks were analysed for each condition.

### 15. Wound-healing assay

The wound-healing assay was performed in Culture-Insert IbiTreat (2 Well 35 mm high μ-Dish; Ibidi). The images were acquired every 30 minutes during 48 hours. The automatic cell and wounded area detection was performed using custom plug-in for ImageJ, based on previously published recommendations (Jonkman *et al*. 2014).

### 16. Transwell^®^ migration and invasion assay

Cells (75×10^3^/passage) were plated into upper compartment of 8 μm pore Transwell^®^ inserts (Falcon™) in a serum-free media, whereas lower compartment was filled with standard growth media, as a chemoattractant.

For invasion assay inserts were pre-coated with 1.25 mg/ml Matrigel™ Basement Membrane Matrix (Becton Dickinson). After 16 hours at 37^0^C, 5 % CO_2_ cells were fixed in ice-cold 100% methanol and stained with 1% crystal-violet in 25% methanol. Inserts were then washed and the upper area of the filter was rubbed dry to eliminate the non-migrated cells. The data acquisition was made using an inverted microscope DMIRE2 (Leica) at ×5 magnification. The results of the cell count from five randomly selected fields were averaged. At least two inserts for each condition were analysed.

### 17. Electrophysiological recordings

Whole-cell transmembrane ion currents were recorded in perforated patch configuration at room temperature. The patch pipette solution supplemented with 100 μg/ml gramicidin was composed of (in mM): 150 KCl, 1 MgCl_2_ and 10 HEPES; pH adjusted to 7.3 with KOH. The cells were bathed in solution containing (in mM): 140 NaCl, 5 KCl, 1 MgCl2, 2 CaCl2, 5 glucose and 10 Hepes; pH adjusted to 7.3 NaOH. Patch pipettes were made from borosilicate glass (WPI) and had free tip resistance 2 - 4 MΩ. The currents were recorded using an Axopatch 200B amplifier (Molecular Devices), and analyzed offline using pClamp (Molecular Devices) and MicroCal Origin (MicroCal Software Inc., Northampton, MA, USA).

### 18. Calcium and sodium imaging

The following solutions were used (in mM): (1) the Hank’s balanced salt solution (HBSS) –NaCl 150, KCl 5, MgCl_2_ 1, CaCl_2_ 2, D-Glucose 10, HEPES 10; pH 7.4 with NaOH; (2) Na^+^-free solution – choline chloride 150, MgCl_2_ 1, KCl 5, CaCl_2_ 2, D-Glucose 5.6, HEPES 10; pH 7.4 with KOH; (3) Ca^2+^-free solution – EGTA-NMDG 5, NaCl 150, MgCl_2_ 3, KCl 5, D-Glucose 5.6, HEPES 10; pH 7.4 with NaOH; (4) high Ca^2+^ solution – NaCl 150, KCl 5, CaCl_2_ 8, D-Glucose 5.6, HEPES 10; pH 7.4 with NaOH.

Cytosolic Ca^2+^ and Na^+^ concentrations were measured using Fura-2-acetoxymethyl ester (AM) (Interchim) and SBFI-AM (Interchim), correspondingly, as previously described (Iamshanova *et al*, 2016). Briefly, the dyes were dissolved in DMSO containing 0.02% Pluronic^®^ F127 and diluted in HBSS to the final concentrations: 1 μM and 7 μM, respectively. The fluorescence was excited with a xenon lamp (300 W) light using an illumination DG4 system (Sutter) equipped with excitation filter pair 340/26 nm and 387/11 nm (wavelength/bandwidth). The fluorescence was acquired with objective 20× using Superfluor Nikon Eclipse Ti-series inverted microscope equipped with the emission filter 510/84 nm and coupled to an EMCCD camera Rolera EM-C2 (Qimaging), and processed using Metafluor 7.7.5.0 software (Molecular Devices).

### 19. Confocal microscopy

Confocal imaging was performed with LSM 510 META confocal workstation using a Plan-Neofluar 40× 1.3 NA or Plan-Apochromat 63× 1.4 NA objectives (Carl Zeiss, Germany). The illumination intensity was attenuated to 0.5-6 % (depending on the laser line) with an acousto-optical tunable filter (Zeiss, Oberkochen, Germany). To optimize signal quality, the pinhole was set to provide a confocal optical section 0.6–2.5 µm, depending on experimental protocol. To avoid any bleed-through of the fluorescence signal in multi-staining experiments, fluorochromes with well separated excitation and emission spectra were used and imaging was performed using the frame-by-frame or line-by-line multitrack mode of the confocal scanner. The photomultiplier gain and offset in each optical channel were set individually to achieve similar signal intensity at each channel and remove sub-signal noise from the images. The adequacy of the imaging protocol applied to the multi-labeled cells was confirmed by control experiments with mono-labeling.

Fluo-4, Mag-Fluo-4, GFP and FM1-43 were excited by 488 nm line of 500 mW Argon ion laser (Laser-Fertigung, Hamburg, Germany) and the fluorescence was captured at wavelengths 505-530 nm or above 505 nm. Alexa Fluor 546, mCherry, Cal-590™, DsRed2 were exited by the 543 nm line of 5 mW Helium/Neon ion laser and the fluorescence was captured at wavelengths above 560 nm. CellMask™ Plsama Membrane Stain was excited by 633 nm line of 15 mW Helium/Neon ion laser and the fluorescence was captured at wavelengths above 650 nm. The MitoSOX™ was excited by 514 nm line of a 500 mW Argon ion laser and the fluorescence was captured at wavelengths above 560 nm. DAPI was excited by 405 nm blue diode laser and the fluorescence was captured at 470-500 nm. Image processing was carried out using LSM 5 software (Zeiss, Oberkochen, Germany) and with custom routines written in IDL (Research Systems, Inc., Boulder, CO, USA). Statistical analysis was performed using MicroCal Origin (MicroCal Software Inc., Northampton, MA, USA).

### 20. Biotinylation

Protein extraction from cell surface fraction was performed with EZ-Link™ Sulfo-NHS-LC-LC-Biotin (Thermo Fisher Scientific).

### 21. Invadopodia fractionation

Protein extraction from invadopodia fraction was performed as described previously (Busco *et al*, 2010).

### 22. Src-family kinase activity assay

After extraction of proteins (see above) without proteinase and phosphatase inhibitors the Src activity was evaluated with ProFluor^®^ Src-Family Kinase Assay (Promega).

### 23. Zymography

Zymography was performed using 1 % gelatin 10% SDS-PAGE. The cells were grown in FBS-free media overnight. The condition media was centrifuged and the protein samples were extracted (see above) without addition of reducing agents and boiling. The zymograms were developed with 0.5 % Coomassie G-250 in 30 % ethanol and 10 % acetic acid.

### 24. Tissue biopsies

Normal prostate tissues (n=58) were obtained from patients without prostate cancer (PCa) underwent cystoprostatectomy for bladder carcinoma.

Hormone naïve clinically localised cancer samples (HNCLC; n=338) were obtained from patients treated with radical prostatectomy for localized PCa.

48 cases of castration resistant prostate cancers (CRPC) were selected from patients treated with exclusive androgen deprivation therapy (ADT). Tissues were collected by transurethral resection, performed due to lower urinary tract symptoms associated with local tumor progression.

Twenty one cases of metastatic prostate cancer were selected from patients with tissues available for analysis: either lymph nodes (n=14) or bone (n=7). Among these, 5 patients (all with bone metastasis) had been previously treated by hormone deprivation but revealed castration resistance.

Written informed consents were obtained from patients in accordance with the requirements of the medical ethic committee.

Prostate adenocarcinoma tissue microarray (PR484) was obtained from US Biomax, Inc.

### 25. Immuno-histochemical analysis

Slides were deparaffinized, rehydrated, and heated in citrate buffer with pH 6 for antigenic retrieval. After blocking for endogenous peroxidase with 3% hydrogen peroxide, the slides were incubated with the primary antibodies (Table S4). Immunohistochemistry was performed using the streptavidin-biotin-peroxydase method with diaminobenzidine as the chromogen (Kit LSAB, Dakocytomation, Glostrup, Denmark). Slides were finally counterstained with haematoxylin. In control experiments corresponding primary antibodies were omitted from the staining protocol.

### 26. *In vivo* xenografts

#### Orthotopic injections

While under isoflurane anaesthesia, 6 weeks old male NMRI Nude Mice (Charles River Laboratories) were injected into the prostate gland with two PC-3 Luc cell lines (shCTL 10 mice, shNALCN 10 mice) suspended in 50 µl PBS. Photons emitted by cancer cells were counted by bioluminescent imaging (FimagerTM; BIOSPACE Lab) and expressed in counts per minute (c.p.m.). Tumor growth was monitored by bioluminescence imaging. Animal weight was measured every week. At necropsy, *ex vivo* BLI measurement for each collected organ was performed within 10 min of D-luciferin intraperitoneal injection (150 mg/kg). Mice were euthanized 12 weeks following implantation of tumor cells and metastatic bioluminescence was measured.

#### Intra-tibial injections

SCID male mice, 6 weeks of age, were housed in barrier conditions under isolated laminar flow hoods. Mice bearing tumor xenografts were closely monitored for established signs of distress and discomfort. Intra-osseous tumor xenograft experiments were performed as previously described (Fradet *et al*, 2013): two PC-3 Luc cell lines (shCTRL 10 mice, shNALCN 10 mice) were suspended as 6 x 10^5^ in 15µL PBS and injected in the bone marrow cavity. Mice were sacrificed after 31 days. Radiographs (LifeRay HM Plus, Ferrania) of animals were taken at that time after inoculation using X-ray (MX-20; Faxitron X-ray Corporation). Hind limbs were collected for histology and histomorphometrics analysis. The bone lesion surface, that includes lytic and osteoblastic regions, was measured using the computerized image analysis system MorphoExpert (Exploranova). The extent of bone lesions for each animal was expressed in mm^2^. Tibiae were scanned using microcomputed tomography (Skyscan1174, Skyscan) with an 8.1 µM voxel size and an X-ray tube (50 kV; 80 µA) with 0.5 µm aluminum filter. Three-dimensional reconstructions and rendering were performed using the manufacturer’s suite (Respectively, NRecon&CTVox, and Skyscan). Bone Volume/Tissue Volume (%BV/TV) includes residual trabecular and remaining cortical bone.

#### Intracardiac injections

5-6 weeks old male NMRI Nude Mice (Charles River Laboratories) were injected into the heart with PC-3 Luc (11 mice) and cells stably overexpressing NALCN (11 mice) suspended in 100 µl PBS. Computed tomography scans were performed on a fast microtomograph Bruker 1278 using an image pixel size of 51.4 µm. The source voltage was 59 kV (753 µA) with a 1 mm aluminium filter. A whole body acquisition consisted in two 360° scans with one projection acquired each 0.5 degrees. Reconstructions were achieved with NRecon software (1.7.1.6, Bruker, Germany) and images were analysed with CTvox software (3.3.0 r1403, Bruker, Germany) and DataViewer software (1.5.6.2, Bruker, Germany) to vizualize respectively, the 3D skeleton volume and the transversal slices of scan areas. During the exam, mice were anesthetized with a mixture consisting of air/isoflurane 2% (Iso-Vet, Piramal Healthcare).

#### PTEN null mice models

PTEN null mice models used in this study were previously described (Parisotto *et al*, 2018). In brief, gene ablation was induced by intraperitoneal injections of Tamoxifen (1 mg/mouse) daily for 5 days to 8 weeks-old mice in order to generate mutant PTEN^(i)pe−/−^ and PTEN/p53^(i)pe−/−^ mice (pe – prostate epithelium, (i) – induced). After Tamoxifen treatment 8 weeks-old mice developed prostatic intraepithelial neoplasia (PIN) in both groups, whereas 72 weeks-old PTEN^(i)pe−/−^ mice developed non-invasive adenocarcinoma and 18 weeks-old PTEN/p53^(i)pe−/−^ mice developed invasive adenocarcinoma. Respective control mice (PTEN^pe+/+^ and PTEN/p53^pe+/+^) were subjected to corresponding Tamoxifen administration and sacrificed at the given age.

### 27. Statistics

*In vitro* data were analyzed using two-tailed Student’s *t*-test to assess the differences between groups. Pairwise comparisons were tested using a non-parametric two-tailed Mann-Whitney *U* test.

Immunohistochemical comparison between tissue biopsies groups was performed using the χ2 test for categorical data and non-parametric Mann-Whitney *U* test and Kruskal-Wallis tests for continuous data.

*In vivo* data for mice studies were compared using non-parametric two-tailed Mann-Whitney *U* test.

In all cases, *P* values <0.05 were considered significant. Statistical analysis was performed using GraphPad Prism 7 (GraphPad Software Inc., San Diego, USA), StatView, version 5.0, (Abacus Concepts, Berkeley, CA) and OriginPro 2015 Beta3 software (1991–2014 OriginLab Corporation). The figures were created using CorelDRAW 11.633 software (2002 Corel Corporation).

### 28. Study approval

Mice for orthotopic injections of prostate cancer cells were bred and housed at the In Vivo platform of the Canceropộle Grand Ouest at Animalerie UTE-IRS1 (Nantes, France) under the animal care license no. C44278. The project was approved by the French national ethical committee (APAFIS #2837-2015112314119496v2). *PTEN* null mice were approved by the Institutional Animal Care and Use Committee of the Emory University (Atlanta, GA), which is accredited by the American Association for the Accreditation of Laboratory Animal Care. Mice for intra-tibial and intracardiac injections were purchased from Charles River and handled according to the French Ministerial Decree No.87-848 of 19 October 1987.

Experimental protocols were approved by the Institutional Animal Care and Use Committee at the Université Lyon-1 (France) (CEEA-55 Comité d’Ethique en Expérimentation Animale DR2014-32) and at CNRS Orléans (CECCOn°3) and received the number 19911 for authorization from the Ministerial Services.

## Acknowledgments

We thank F. Cabon (Cancer Research Centre of Toulouse, France) and S. Fraser (Imperial College London, United Kingdom) for C4-2 and PC-3M cell lines, accordingly. We thank E. Dewailly (Inserm U1003, University of Lille, France) for help with reagents and cellular biology methods. We thank C. Slomianny, E. Richard and A.-S. Lacoste of the BICeL-Campus Scientific City Facility for access to instruments and technical advices. We are indebted to the Research Federation FRABio (University of Lille, CNRS, FR 3688, FRABio, Biochimie Structurale et Fonctionnelle des Assemblages Biomoléculaires) for providing the scientific and technical environment conducive to achieving this work. We thank Dr. C. Lemmers (Vectorology facility, PVM, Biocampus Montpellier, CNRS UMS3426) for lenvirus production. We thank C. Lagadec and R.-A. Toillon for help with their flow-cytometry facility (Inserm U908, University of Lille, France). We thank Daniel Metzger (Inserm U1258, University of Strasbourg, France) and Emmanuelle Germain (Inserm U1003, University of Lille, France) for providing us with cDNA from *PTEN* null mice model. We also thank Pr. Stephan Reshkin (University of Bari) and Pr. Philippe Chavrier (Curie Institute, Paris) for helpful discussions.

## Author contributions

O.I., D.G., Ar.M. and N.P. conceived and planned the study. O.I. and A.F. designed, performed and analyzed cell culture assays (e.g. cell cycle analysis, cellular viability, apoptosis, proliferation, cellular motility, wound-healing, Transwell^®^ migration and invasion assays) as well as invadopodia fractionation and Src-family kinase activity assay. D.G. designed, performed and analyzed confocal microscopy experiments. O.I., A.F., R.D., E.D. and A.B. designed, performed and analyzed cellular and molecular biology experiments (e.g. RT-qPCR, immunoblotting, biotinylation, zymography). G.S., P.M. and A.F. designed, performed and analyzed electrophysiological recordings. O.I., A.F. and P.M. designed, performed and analyzed Ca^2+^ and Na^+^ imaging experiments. Ar.M. constructed NALCN-expressing plasmids. V.L. participated in choosing the epitope for antibody against NALCN. A.H. participated in the screening of shNALCN clones. L.A. and A.B. amplified and extracted plasmid constructs. H.I. and Ar.M. established PC-3 cell line stably overexpressing NALCN. C.S. designed, performed and analyzed time-lapse video microscopy, cellular motility and wound-healing experiments. L.B. and Se.R. designed, performed and analyzed measurements of intracellular pH. T.O. and S.M.-L. designed, performed and analyzed tumor growth and metastases in mice after orthotopical injections of PC-3 cells. Se.R., S.C., St.R., M.L., J.S. and S.L. designed, performed and analysed the bone metastatic model in mice after intracardiac injections of PC-3 cells. S.G. and P.C. designed, performed and analyzed osteoblasts-mediated invasion of PC-3 cells, intratibial injections of the latter and analysis of produced bone lesions. Ad.M. and P.G. performed immunohistochemical staining of prostate adenocarcinoma tissue microarray and V.L. helped them. G.F.-H. performed immunohistochemical staining of prostate biopsies from patients and analyzed the correlation between expression of different proteins. O.I., D.G., and N.P. wrote the manuscript and O.I., D.G., A.F., M.D., Se.R., Ar.M., P.C. and G.F.-H. reviewed it.

## The paper explained

Our study is focused on molecular identification of Na+/Ca2+ signaling pathway driving metastatic behavior of prostate cancer cells. Indeed, metastasis is the main cause of prostate cancer patient’s mortality. It is, therefore, vital to understand the processes underlying the progression of prostate cancer to aggressive metastatic phenotypes. Alteration of ionic homeostasis driven by aberrant ion channel expression/function, referred to as ‘oncochannelopathies’ (Prevarskaya et al., Physiological Reviews 2018), is one of the important features of the development of invasive tumor phenotype. However, the causal link between aberrant ionic homeostasis and invasion of cancer cells (in particular in prostate cancer) as well as the underlying molecular mechanisms remain poorly understood.

Using variety of in vitro and in vivo approaches, we demonstrate that strongly metastatic cancer cells acquire specific ionic signature that facilitates persistent invasion. In particular, we show how Na+ influx via the leak channel, NALCN, initially identified in neurons, gives rise to long-lasting Ca2+ oscillations (in cytoplasm, endoplasmic reticulum and mitochondria), ROS production and promotes invasion.

We also show that modulation of NALCN bioavailability strongly affects prostate cancer progression in vivo. As our study provides an example of application and combination of innovative experimental approaches at multiple levels of assessment allowing uncovering the fundamental mechanisms of carcinogenesis.

## Availability for supporting data

We declare that the material of the present manuscript has been neither published nor submitted for publication elsewhere, in part or in whole. All data generated or analysed during this study are included in this published article (and its supplementary information files).

## Additional table and figures legends

**Table S1.** NALCN expression in hormone naïve clinically localized cancer.

| NALCN |  | - | + | ++ | P |
| --- | --- | --- | --- | --- | --- |
| ISUP group | 1 | n = 47 | n = 25 | n = 3 | 0.0003 |
|  | 2 | n = 39 | n = 42 | n = 10 |  |
|  | 3 | n = 37 | n = 73 | n = 17 |  |
|  | 4 | n = 10 | n = 9 | n = 0 |  |
| Stage | pT2 | n = 97 | n = 93 | n = 10 | 0.0003 |
|  | pT3 | n = 36 | n = 56 | n = 20 |  |

**Figure S1.**
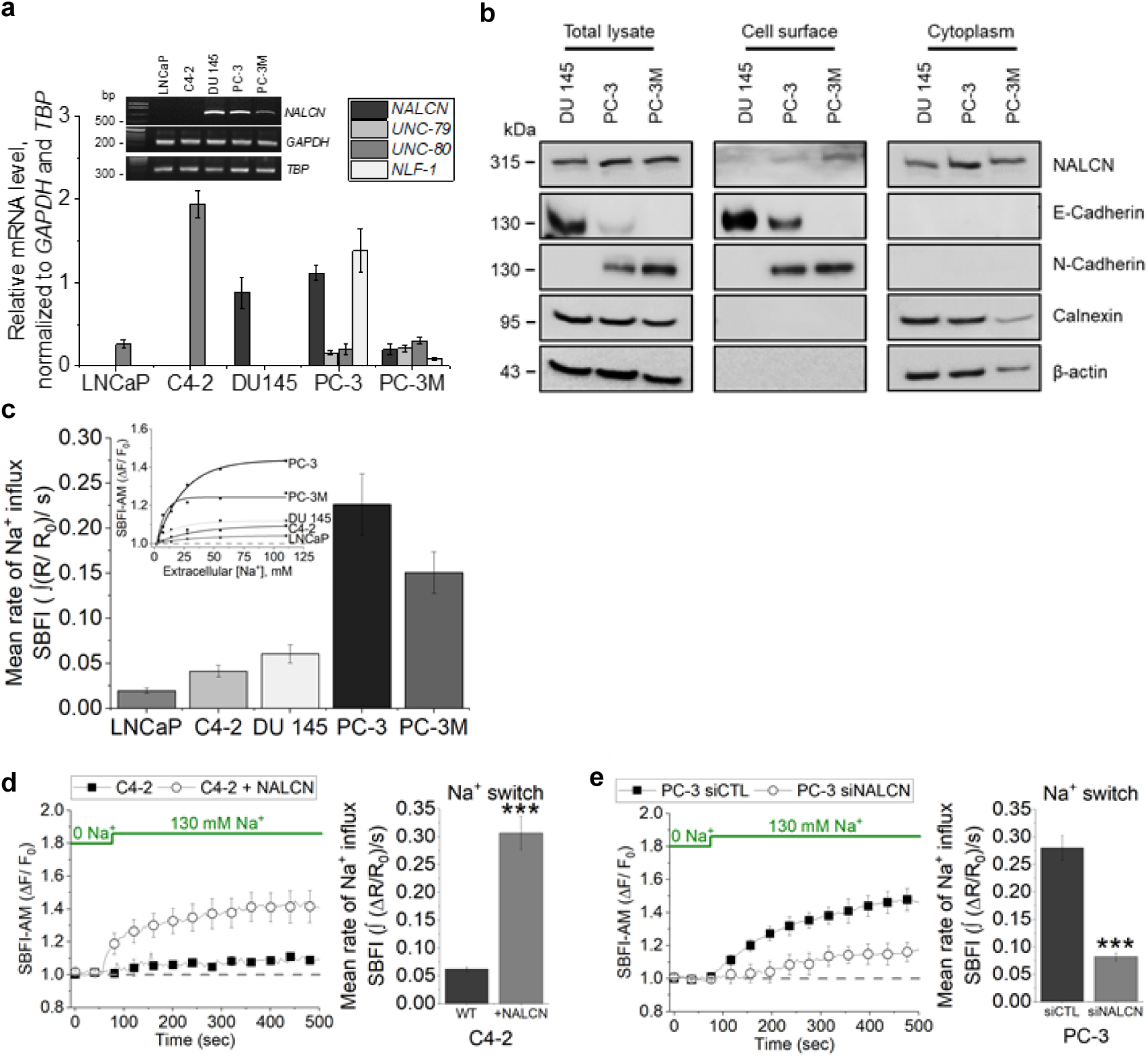
Human prostate cancer cells express functional NALCN that mediates Na^+^ influx. (**a**) Conventional and RT-qPCR for NALCN channelosome genes in human prostate cancer (PCa) cell lines. Data are normalised to GAPDH and TBP and are presented as mean values±SEM of five independent experiments (n=5). (**b**) NALCN was found in plasma membrane biotinylated fraction of human prostate cancer cells endogenously expressing the channel. E-Cadherin and N-Cadherin were used as plasmalemmal markers, whereas calnexin and β-actin – as markers of cytoplasmic fraction. (**c**) Background Na^+^ influx measured in human prostate cancer cell lines using ratiometric dye SBFI-AM. The bar diagram plots: mean signal temporal densities per cell (n=40-60). (**d**) In weakly metastatic C4-2 cells basal Na^+^ influx was induced by transient NALCN overexpression (+NALCN), when compared with wild-type cells (WT). The bar diagram plots: mean signal temporal densities per cell (n=27-62). (**e**) In highly metastatic PC-3 cells transient NALCN silencing with siRNA (siNALCN) resulted in suppression of basal Na+ influx, when compared with siRNA targeting firefly luciferase (siCTL). The bar diagram plots: mean signal temporal densities per cell (n=203-241). In (**d**) and (**e**) cells were transfected for 72 hours. Data are presented as mean values±SEM of three independent experiments. ***P<0.001, two-tailed Student’s *t*-test.

**Figure S2.**
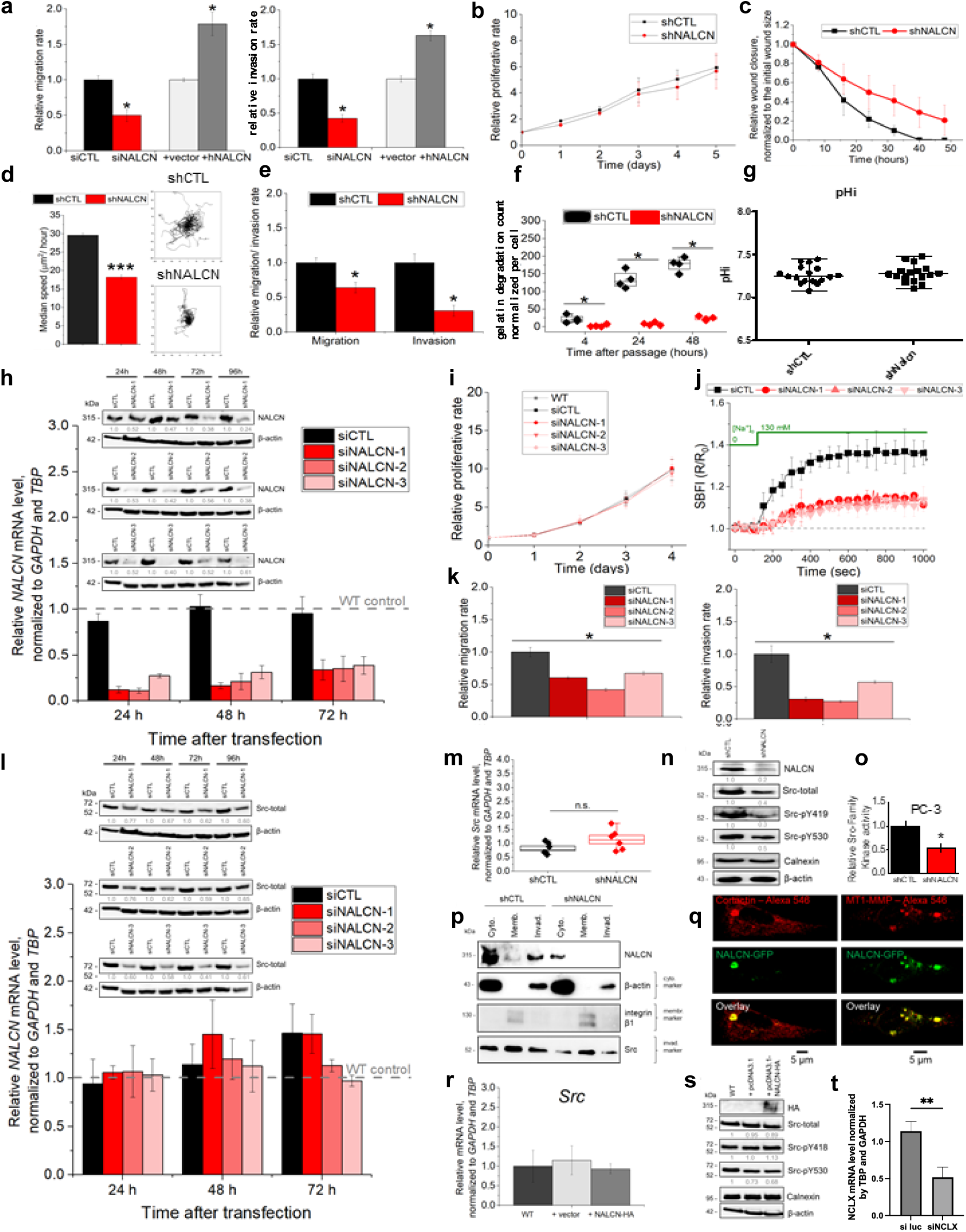
Role of NALCN in metastatic hallmarks of PC-3 cells. (**a**) Transwell® migration and Matrigel invasion assays after transient NALCN suppression and overexpression. Control cells were transfected for 72 hours whether with siRNA targeting firefly luciferase (siCTL) or with an empty vector pcDNA3.1 (+vector), whereas NALCN depleted cells – with siRNA-1 (siNALCN) and NALCN overexpressing cells with pcDNA3.1-NALCN-HA (+NALCN). Data are presented as mean±S.E.M of four independent experiments. *P<0.05, two-tailed Mann-Whitney U test. (**b**) Cellular proliferation assessed by sulforhodamine B assay. (**c**) Wound-healing was slowed down in NALCN depleted cells. (**d**) Cellular motility (calculated as μm2/hour) was remarkably inhibited in NALCN suppressed cells. Data are mean±S.E.M (n=838-1629). ***P<0.001, two-tailed Student’s t-test. Motility traces of individual control (shCTL) and NALCN downregulated (shNALCN) cells (n=30). (**e**) Transwell® migration and Matrigel invasion assays for cells with stable NALCN downregulation. Data are mean±S.E.M of four independent experiments. *P<0.05, two-tailed Mann-Whitney U test. (**f**) Count of gelatin degradation dots produced by control (shCTL) and stable NALCN knockdown (shNALCN) cells. Box-plots with raw data from four independent experiments are shown (n=4). *P<0.05, two-tailed Mann-Whitney U test. (**g**) Intracellular pH was not affected by NALCN knockdown (n=17). (**h**) NALCN mRNA and protein levels in non-transfected (WT control) cells and after transfection with siRNA targeting firefly luciferase (siCTL) or transient NALCN silencing with siRNA-1 (siNALCN-1), siRNA-2 (siNALCN-2), and siRNA-3 (siNALCN-3). RT-qPCR data are normalised to GAPDH and TBP. Immunoblotting data are normalised to β-actin. Regardless of siRNA applied transient NALCN silencing did not affect proliferation (**i**) of PC-3 cells, while suppressing the basal Na+ influx (**j**). Na+ imaging data are mean values±S.E.M. of three independent experiments per cell (n=203-241). (**k**) Transwell® migration and Matrigel invasion assays as in (**a**). Data are mean±S.E.M of four independent experiments (n=4). *P<0.05, two-tailed Mann-Whitney U test. (**l**) Src mRNA and protein levels after transfection with siRNA targeting firefly luciferase (siCTL) or transient NALCN silencing with siRNA-1 (siNALCN-1), siRNA-2 (siNALCN-2), and siRNA-3 (siNALCN-3). Src total– total protein level, Src-pY419 – active form of Src, Src-pY530 – nonactive form of Src. RT-qPCR data are normalised to GAPDH and TBP. Immunoblotting data are normalised to β-actin. (**m**) RT-qPCR for Src in control (shCTL) and cells with stable NALCN suppression (shNALCN). Data are normalised to GAPDH and TBP and are presented as mean±SEM of six independent experiments (n=6). n.s. – non-significant. (**n**) Representative immunoblot with band intensities calculated for Src protein level as in (**m**). Data are normalised to β-actin and calnexin. (**o**) Kinase activity assessed with ProFluor® Src-Family kinase assay. Data are mean values±S.E.M of four independent experiments (n=4). *P<0.05, two-tailed Mann-Whitney U test. (**p**) Representative immunoblot of NALCN localisation in cytoplasmic, membrane and invadopodial fractions after stable NALCN knockdown. (**q**) Representative confocal images of co-localisation between green fluorescent protein (GFP)-tagged NALCN and invadopodial markers: cortactin and MT1-MMP. (**r**) RT-qPCR for Src after transient overexpression of haemagglutinin (HA)-tagged NALCN (+NALCN-HA). Data are normalised to GAPDH and TBP. (**s**) Representative immunoblots with band intensities obtained for Src protein level as in (**r**). Data are normalised to β-actin and calnexin. (**t**) RT-qPCR for NCLX in control (siLuc) and cells transfected by siNCLX. In (**b**), (**c**), (**h**), (**i**), (**l**), (**n**), (**r**), (**s**) and (**t**) data are presented as mean values±S.E.M of three independent experiments.

**Figure S3.**
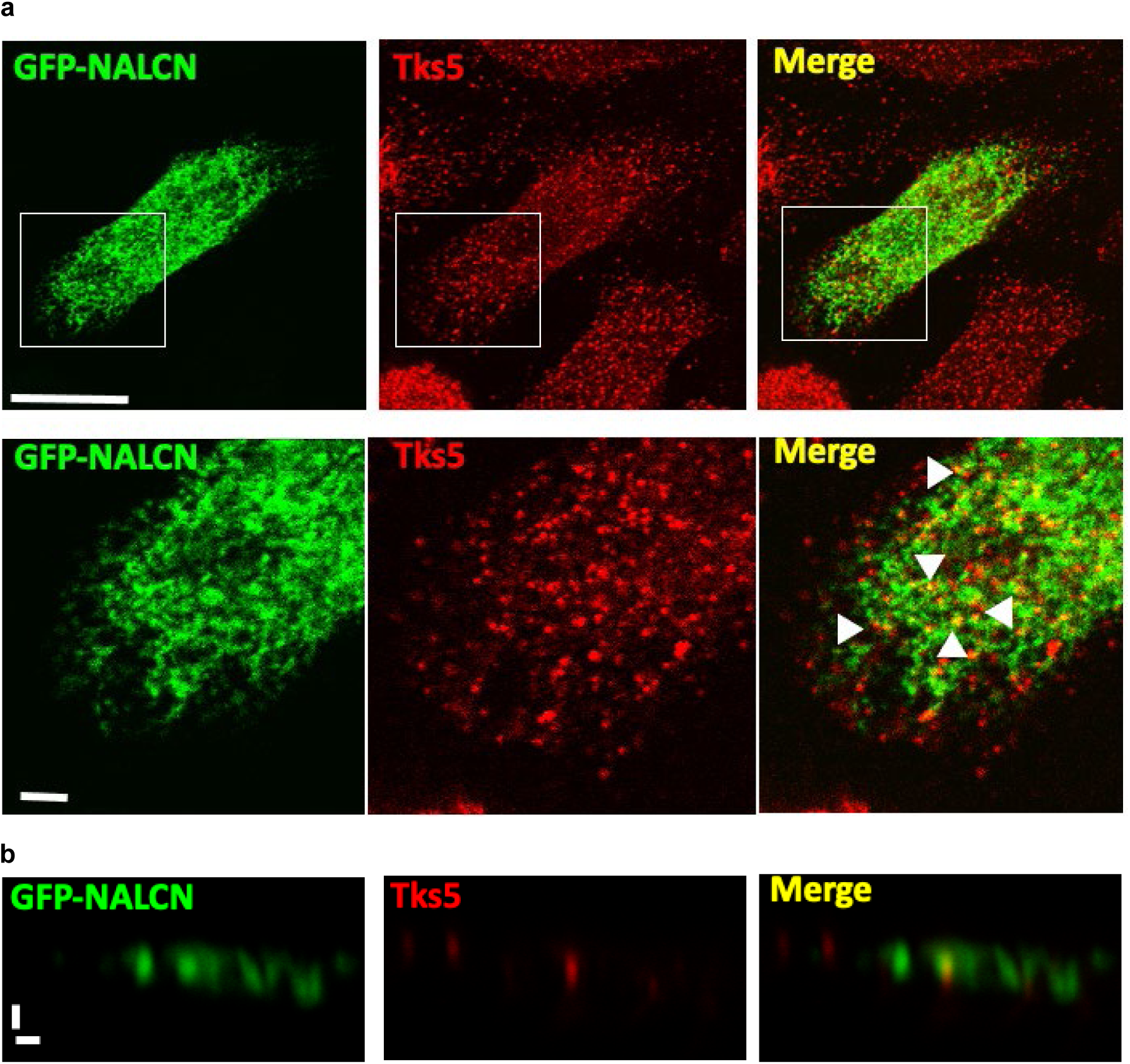
Invadosome formation on gelatin layer. (**a**) Confocal microscopy images of PC3 cells plated on gelatin and transfected with NALCN GFP, treated with 10% FBS in RPM1 overnight, and stained for endogenous Tks5 (in red) to visualize invadosomes. Scale bar, 15 μm. Lower inset panels highlight the colocalization of NALCN and Tks5. Scale bar, 1 μm. (**b**) Confocal microscopy XZ section of PC3 cells plated on gelatin and transfected with NALCN GFP treated with 10% RPM1 overnight and stained for endogenous Tks5 (in red) to visualize invadosomes. Scale bar, 1 μm.

**Figure S4.**
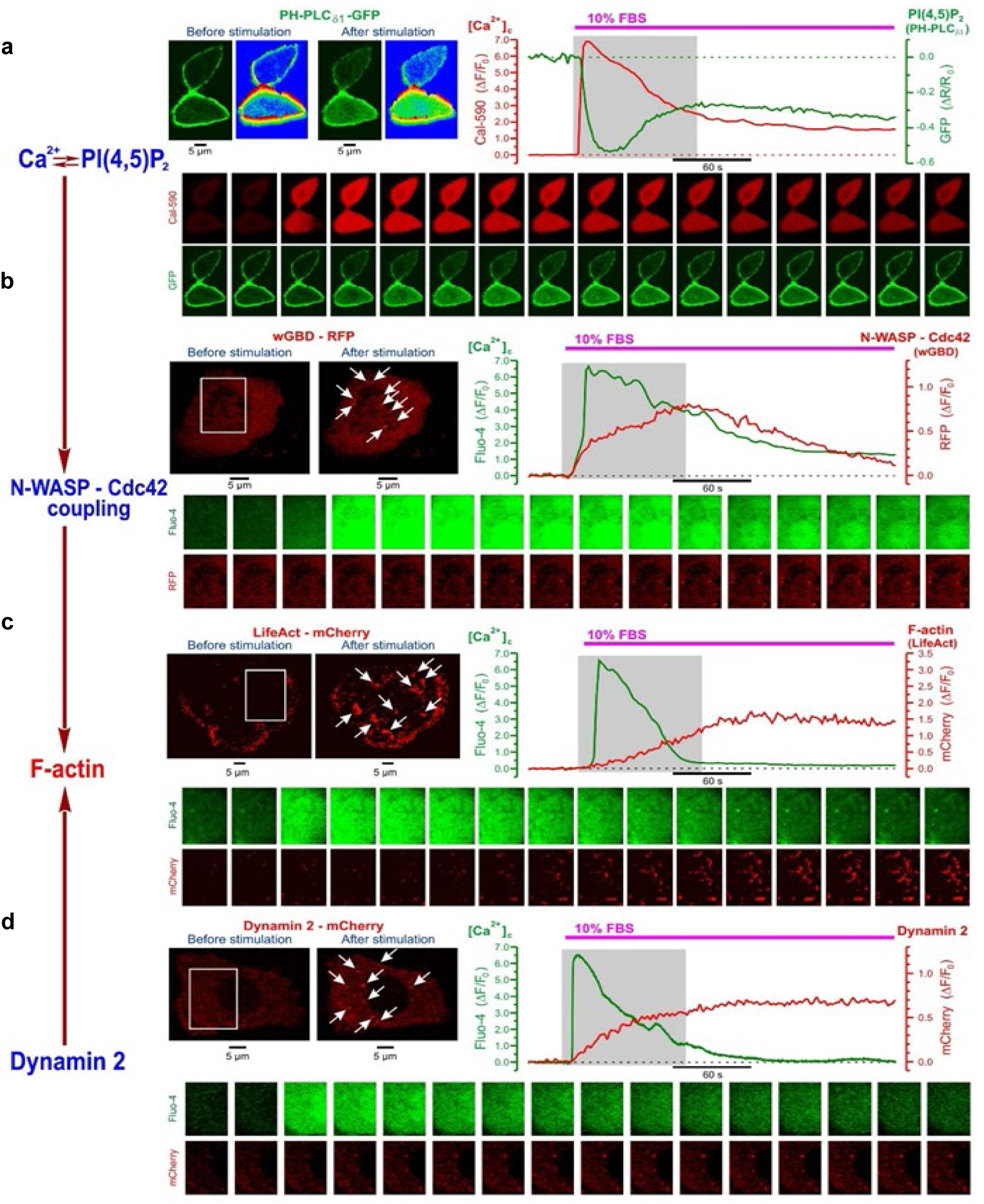
FBS-induced [Ca^2+^]c response in PC-3 cells is associated with changes in PI(4,5)P2, N-WASP/Cdc42, dynamin and actin consistent with initiation of invadopodia formation. (a) Left: confocal images of PH-PLC_∂1_ - GFP fluorescence captured before and 1 min after stimulation with 10% FBS, and their rainbow-coded 3D shaded-surface plots. Right: corresponding traces of relative (R=Fplasmalemma /Fcytosol) changes in GFP fluorescence (green), reflecting translocation PH-PLC_∂1_ from the plasma membrane to cytosol caused by PI(4,5)P2 degradation, and in Cal-590 fluorescence (red) reflecting [Ca^2+^]c dynamics. (b) Left: near cell bottom confocal (< 0.8 µm) images of wGBD - RFP fluorescence before and 2 min after stimulation with 10% FBS. Note puncta formation (arrows) reflecting spots where Cdc42 is activated to bind N-WASP. Right: corresponding traces of relative changes in RFP fluorescence (red), reflecting the dynamics of Cdc42 activation, and in Fluo-4 fluorescence (green) reflecting [Ca^2+^]c dynamics. (c) Left: near cell bottom confocal (< 0.8 µm) images of PC-3 cell transfected with F-actin marker LifeAct - mCherry were captured before and 3 min after stimulation with 10% FBS. Note F-actin-enriched structures (arrows) induced by stimulation with FBS. Right: corresponding traces of relative changes in mCherry fluorescence (red), reflecting F-actin dynamics, and in Fluo-4 fluorescence (green) reflecting [Ca^2+]^c dynamics. (d) Left: near cell bottom confocal (< 0.8 µm) images of PC-3 cell transfected with Dynamin 2 - mCherry were captured before and 2 min after stimulation with 10% FBS. Note dynamin puncta (arrows) induced by stimulation with FBS. Right: corresponding traces of relative changes in mCherry fluorescence (red) reflecting the dynamics of the dynamin puncta formation and in Fluo-4 fluorescence (green) reflecting [Ca^2+^]c dynamics. The galleries below the plots show images (a) or enlarged boxed regions of the images (b-d) captured during the periods highlighted on the plots by grey background: every 3rd image (a-c), every 35th image (d).

**Figure S5.**
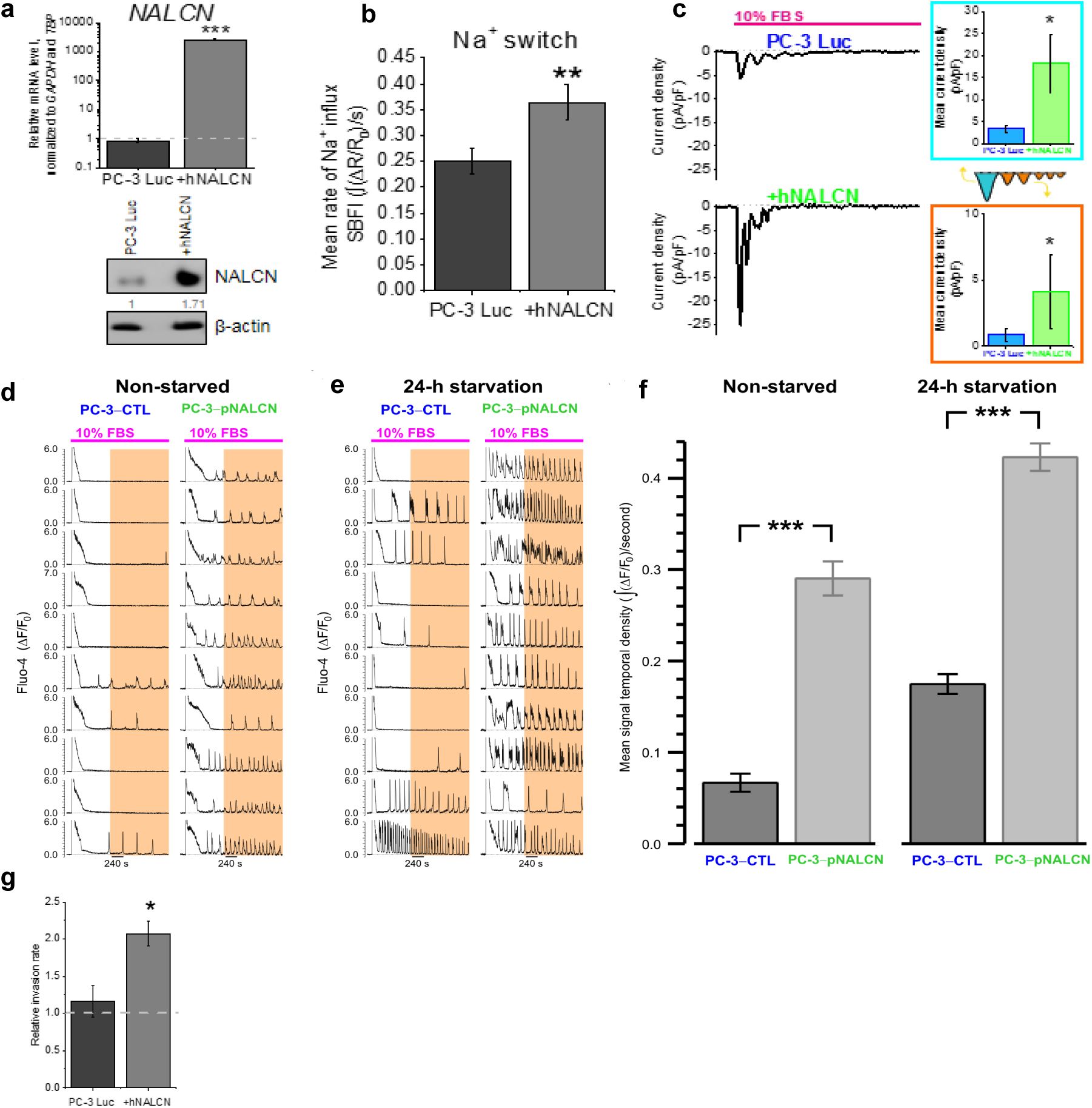
NALCN overexpression promotes metastatic cell behaviors in vitro and in vivo. (**a**) NALCN mRNA and protein levels in control human prostate cancer cells permanently overexpressing luciferase (PC-3 Luc) and in cells stably overexpressing NALCN (+hNALCN). RT-qPCR data are normalised to GAPDH and TBP and are presented as mean values±S.E.M of three independent experiments (n=3); and related to mRNA level of PC-3 stably overexpressing mCherry (grey dashed line). ***P<0.001, two-tailed Mann-Whitney U test. Immunoblotting data are normalised to β-actin and are presented as mean of three independent experiments (n=3). (**b**) Mean rate of Na^+^ (SBFI) influx calculated as signal mass (∫(ΔR/R0)/s) in PC-3 Luc and +hNALCN cells. The bar diagram plots: mean signal temporal densities per cell (n=141-287). Data are presented as mean ±S.E.M. of three independent experiments. **P<0.01, two-tailed Student’s t-test. (**c**) Representative traces of the inward Na^+^ current recorded using perforated patch-clamp technique. The FBS-induced inward Na^+^ current in PC-3 was significantly larger in cells stably overexpressing NALCN than in control cells. The bar diagram plots: mean current densities (pA/pF; mean±S.E.M.) per cell (n=4-6) during initial transient (cyan) and oscillations (orange). *P<0.05, two-tailed Mann-Whitney U test. (**d**) and (**e**) Representative traces of FBS-induced [Ca^2+^]c responses reported by confocal time-series imaging of fluo-4 fluorescence. Overexpression of NALCN facilitates [Ca^2+^]c oscillations in non-starved and pre-starved PC-3 cells. (**f**) Mean signal (Fluo-4) temporal densities during highlighted periods were compared between PC-3 Luc and +hNALCN in non-starved (n=259 and n=254, respectively) and pre-starved (n=360 and n=317, respectively) cells. Data are mean values±S.E.M. ***P<0.001, two-tailed Student’s *t*-test. (**g**) NALCN overexpression significantly increases invasiveness of PC-3 cells as reported by Transwell® Matrigel invasion assay. Data are presented as mean±S.E.M of four independent experiments and related to invasion rate of PC-3 stably overexpressing mCherry (grey dashed line). *P<0.05, two-tailed Mann-Whitney U test.

**Figure S6.**
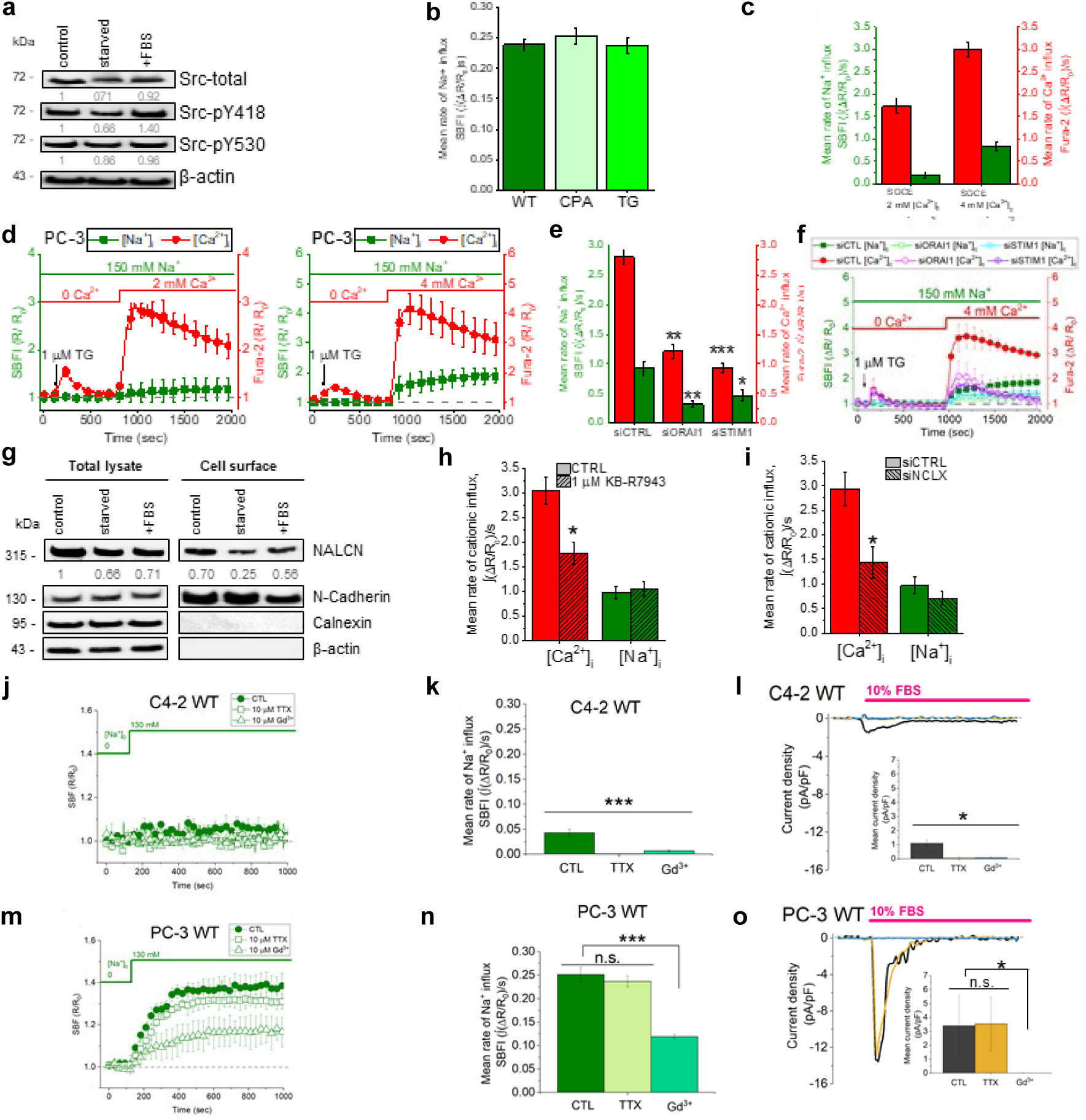
Interplay between store-operated Ca^2+^ entry (SOCE) and NALCN in PC3 cells. (**a**) Representative immunoblot with band intensities calculated for Src protein level in control, 24 h prestarved cells and FBS-stimulated cells after starvation (+FBS). Src total– total protein level, Src-pY419 – active form of Src, Src-pY530 – nonactive form of Src. Data are normalised to β-actin and are presented as mean values of three independent experiments (n=3). (**b**) Mean rate of Na^+^ influx calculated as SBFI-AM signal mass (∫(ΔR/R0)/s) at different concentrations of extracellular Ca^2+^ ([Ca^2+^]o) and after ER depletion with 50 μM cyclopiazonic acid (CPA) and 1 μM thapsigargin (TG). The bar diagram plots: mean signal temporal densities per cell (n=45-360). (**c**) Mean rate of Na^+^ (SBFI) and Ca^2+^ (Fura-2) influx calculated as signal mass (∫(ΔR/R0)/s) during different SOCE components. The bar diagram plots: mean signal temporal densities per cell (n=283-360). (**d**) SOCE-induced changes of cytosolic concentrations of Na^+^ ([Na^+^]c, SBFI) and Ca^2+^ ([Ca^2+^]c, Fura-2) using 2 mM [Ca^2+^]o and 4 mM [Ca^2+^]o. (**e**) and (**f**) SOCE-induced changes of [Na^+^]c (SBFI) and [Ca^2+^]c (Fura-2) in cells transfected for 48 hours with siRNA targeting firefly luciferase (siCTL), ORAI1 (siORAI1) and STIM1 (siSTIM1). Mean rate of Na^+^ (SBFI) and Ca^2+^ (Fura-2) influx calculated as signal mass (∫(ΔR/R0)/s) after silencing of major SOCE players. The bar diagram plots: mean signal temporal densities per cell (n=109-182). *P<0.05, **P<0.01, ***P<0.001, two-tailed Student’s t-test. (**g**) Representative immunoblot with band intensities calculated for NALCN protein level in total cell lysate and biotinylated plasma membrane fraction. N-Cadherin was used as plasmalemmal marker, whereas ER-residing protein calnexin and cytoskeletal β-actin – as markers of cytoplasmic fraction. For cytoplasmic fraction data are normalised to β-actin and calnexin, whereas for plasmalemmal fraction data are normalised to N-Cadherin, all data are presented as mean values of three independent experiments (n=3). (**h**) and (**i**) Mean rate of Na^+^ (SBFI) and Ca^2+^ (Fura-2) influx calculated as signal mass (∫(ΔR/R0)/s) after pharmacological inhibition of NCX-reverse mode (1 μM KB-R7943) (h) and mitochondrial NCLX silencing (siNCLX) (**i**). The bar diagram plots: mean signal temporal densities per cell (n=134-289). *P<0.05, two-tailed Student’s *t*-test. (**j**) and (**m**) Background Na^+^ influx measured using ratiometric dye SBFI-AM in NALCN-negative human prostate cancer cell line, C4-2 WT, and endogenously expressing NALCN human prostate cancer cell line, PC-3 WT. In the latter, Na^+^ influx is tetrodotoxin (TTX)-resistant and gadolinium (Gd3+)-sensitive, as previously described for NALCN-mediated current (INALCN) in neurons. (**k**) and (**n**) The bar diagram plots: mean signal temporal densities per cell (for C4-2, n=702-772; for PC-3, n=1117-1571). Data are presented as mean ±S.E.M. of six independent experiments. n.s. – non-significant, ***P<0.001, two-tailed Student’s *t*-test. (**i**) and (**o**) Representative traces of the inward Na^+^ current recorded using perforated patch-clamp technique. In C4-2 cell line, TTX or Gd^3+^ applications both completely abolished relatively small FBS-induced inward Na^+^ current. In contrast, the FBS-induced inward Na^+^ current in PC-3 cells was significantly larger, resistant to TTX but sensitive to Gd^3+^. The bar diagram plots: mean current densities (pA/pF; mean±S.E.M.) per cell (n=4-6). n.s. – non-significant, *P<0.05, two-tailed Mann-Whitney U test. In (**b**), (**c**), (**d**), (**f**), (**i**) and (**j**) data are presented as mean values±SEM of three independent experiments.

**Figure S7.**
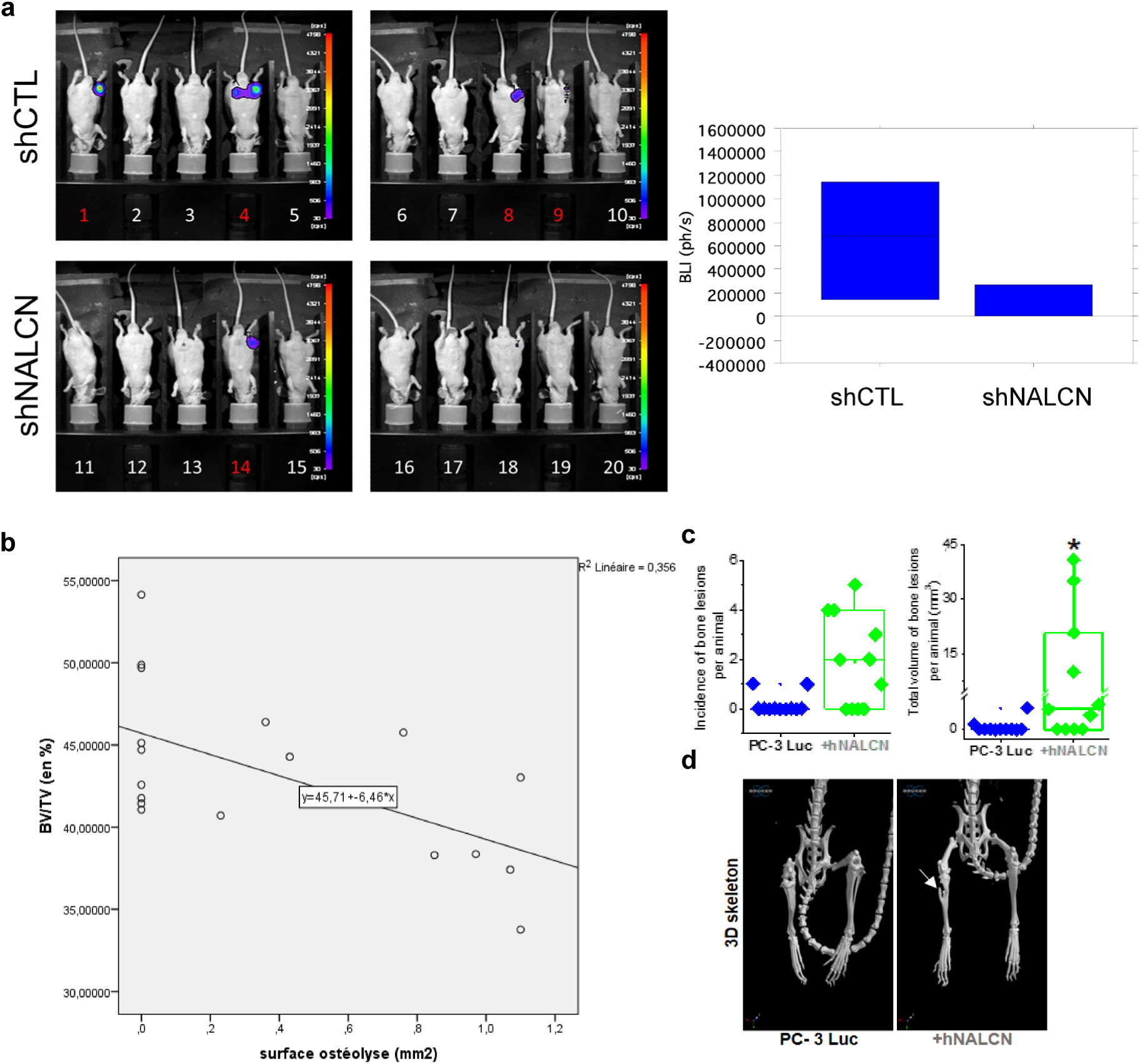
Analysis of bone tissue destruction. (**a**) Bioluminescence of mice 31 days after intra-tibial injection of control (PC-3 Luc-shCTL) or NALCN depleted cells (PC-3 Luc-shNALCN) with corresponding signal analysis (photons/s). Data are mean±S.E.M. (**b**) Correlation between osteolysis surface (mm^2^) and bone volume/total volume (BV/TV, %). P =0,009. two-tailed Mann-Whitney U test. (**c**) Incidence and total volume of bone lesions in mice 8 weeks after intracardiac injections with control (PC-3 Luc) and cells stably overexpressing NALCN (+hNALCN). Box-plots with raw data from each mouse are shown. *P<0.05, two-tailed Mann-Whitney U test. (**d**) 3D illustrations of representative skeletal lesions.

**Figure S8.**
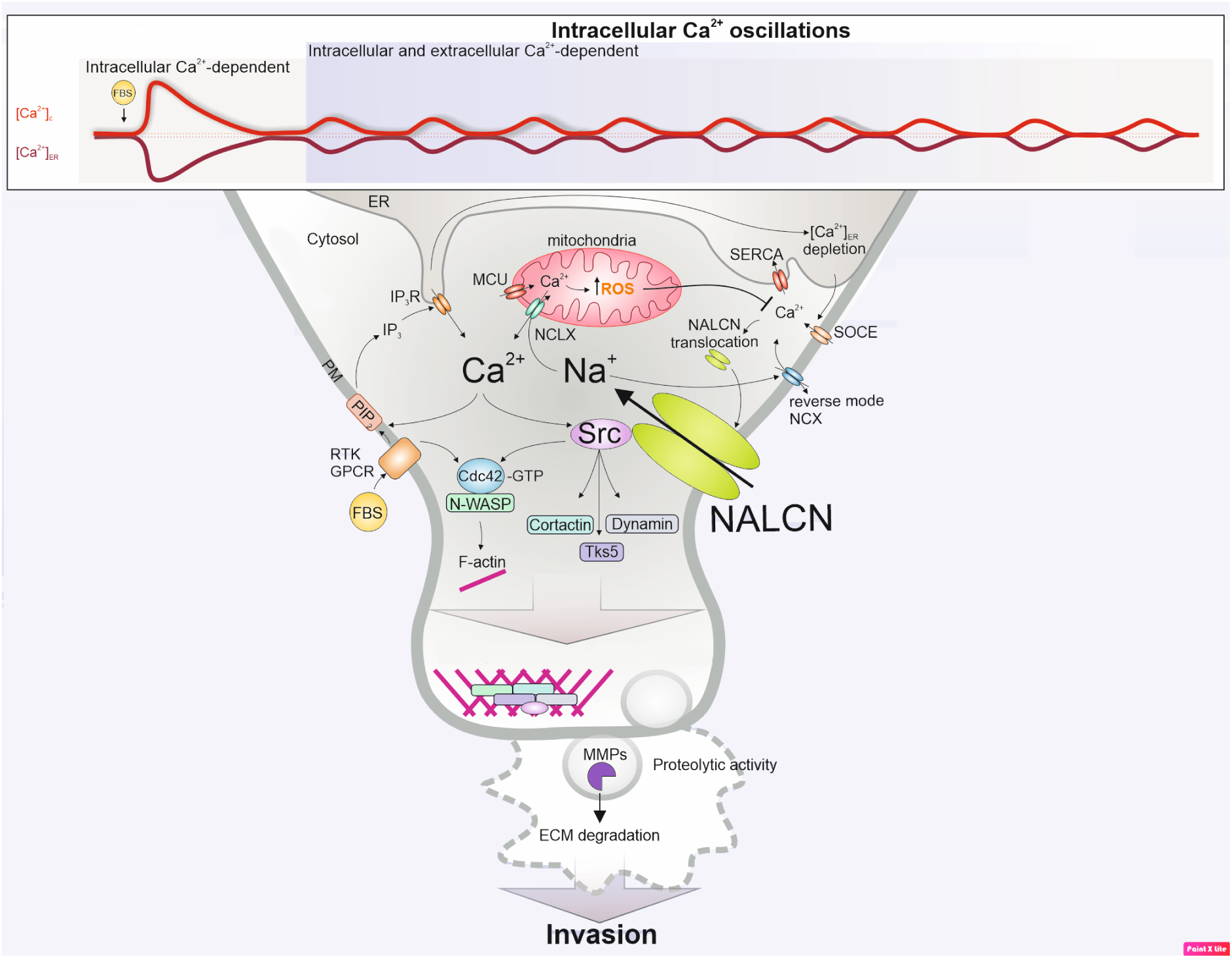
Schematic representation of signaling pathways employed by NALCN to promote cancer cell invasiveness. Abbreviations: PM – plasma membrane, ER – endoplasmic reticulum, NALCN – Na^+^ leak channel non-selective, RM-NCX – reverse mode of Na^+^/ Ca^2+^ exchanger, NCLX – mitochondrial Na^+^/ Ca^2+^ exchanger, IP3R – inositol 1,4,5-trisphosphate receptor, SERCA –sarco/endoplasmic reticulum Ca^2+^-ATPase, MCU – mitochondrial Ca^2+^ uniporter, RTK – receptor tyrosine kinases, GPCR – G-protein coupled receptors, Src – proto-oncogene Src tyrosine kinase, N-WASP – neural Wiskott-Aldrich syndrome protein, ECM – extracellular matrix, MMPs – matrix metalloproteinases, GTP – guanosine-5’-triphosphate, ROS – reactive oxygen species, [Ca^2+^]c – cytosolic Ca^2+^ concentration, [Ca^2+^]ER – Ca^2+^ concentration in ER, [Ca^2+^]mito – Ca^2+^ concentration in mitochondria, [Na^+^]c – cytosolic Na^+^ concentration, [ROS]mito – ROS concentration in mitochondria. Briefly: (i) IP3R-mediated Ca^2+^ release caused by FBS-induced activation of RTK/GPCR causes the ER Ca^2+^ depletion and SOCE; elevation of [Ca^2+^]c in sub-PM microdomains facilitates NALCN translocation to PM and Na^+^ influx promoting RM-NCX and additional Ca^2+^ influx; this triggers SERCA-mediated Ca^2+^ uptake into the ER and MCU-mediated Ca^2+^ uptake into mitochondria; the latter facilitates production of ROS, known to inhibit SERCA, RM-NCX and SOCE elements, and is opposed by NCLX exchanging Na^+^, delivered by NALCN, to mitochondrial Ca^2+^; these positive and negative feedbacks give rise to [Ca^2+^]c oscillations maintaining the activity Src, known as essential component of NALCN channelosome; active Src phosphorylates downstream proteins (cortactin, dynamin and Tks5) recruiting them to actin polymerization regions and giving rise to “invadopodia puncta”. (ii) RTK/GPCR activation is linked to activation of the Rho family GTPase, Cdc42, leading to its binding with N-WASP and subsequent actin nucleation. (iii) Invadopodia maturation is facilitated by Ca^2+^-dependent secretion of MMPs degrading ECM.

